# PFOS aggravates atherosclerosis via *Bacteroides caecimuris* expansion-driven bile acid remodeling and subsequent intestinal FXR–TLR3 signaling cascade

**DOI:** 10.64898/2026.07.01.735947

**Authors:** Lehua Jiang, Shan Huang, Zhen Xu, Rongxin Guo, Jiahui Zhu, Haowei Liang, Chao Yuan, Zhenghao Zhao, Fengyuan Lv, Yutong Ai, Kun Xu, Yutong Wu, Xiaosong Li, Guoqiang Qin, Chenyu Li, Shengqi Hu, Tong Liu, Mengqian Zhang, Zhixuan Zhou, Yang Li, Beining Liu, Qing Wu, Kangyin Chen, Zhongze Fang

## Abstract

**BACKGROUND:** Perfluorooctane sulfonate (PFOS) is a widely distributed persistent organic pollutant in the environment and has been associated with an increased risk of atherosclerosis. However, the underlying pathogenic mechanisms remain largely unclear. This study aimed to investigate the effects of PFOS on atherosclerosis and its associated gut–vascular axis.

**METHODS:** Pseudo–germ-free mouse models and fecal microbiota transplantation (FMT) were used to determine the role of the gut microbiota in PFOS-induced atherosclerosis. Metagenomic sequencing was performed to characterize alterations in gut microbial composition following PFOS exposure, and targeted metabolomics was used to assess bile acid profiles in the ileum and plasma. Transcriptomic analysis of *Bacteroides caecimuris* (*B.caecimuris*) was conducted to explore the reasons for the increased abundance of *B.caecimuris* after PFOS exposure. In addition, intestinal transcriptomics and ChIP-qPCR were performed to validate transcriptional regulation within the FXR–TLR3 signaling axis.

**RESULTS:** Among 127 participants with paired serum and fecal samples, including 82 patients undergoing coronary angiography with Gensini scores (GS score), fecal PFOS levels were significantly associated with lipid profiles and GS score, whereas serum PFOS showed no such association.Mechanistically, PFOS exposure promotes intestinal enrichment of *B.caecimuris* by upregulating its *tolC* gene, thereby enhancing efflux capacity.This microbial shift was accompanied by reduced levels of tauro-ursodeoxycholic acid (TUDCA) and aberrant activation of intestinal FXR signaling.Further analyses demonstrated that FXR activation upregulated TLR3 expression and promoted inflammatory responses and atherosclerosis progression via the TLR3–NF-κB signaling axis. Both intestinal epithelial-specific FXR deficiency (*Fxr*^ΔIE^) and TUDCA supplementation significantly suppressed pathway activation and alleviated disease phenotypes.Functional experiments identified TLR3 as a key downstream effector of FXR. Overexpression of TLR3 abolished the protective effects observed in *Fxr*^ΔIE^ mice. Moreover, pharmacological inhibition of TLR3 using CU CPT-4a significantly improved established atherosclerotic lesions in vivo.

**CONCLUSIONS:** This study identifies a gut microbiota–driven FXR–TLR3 signaling axis that mediates PFOS-induced atherosclerosis. These findings provide new mechanistic insights into environmentally induced cardiovascular disease and suggest potential targets for risk assessment and therapeutic intervention.

## Introduction

Cardiovascular disease remains a leading cause of mortality and disease burden worldwide^[1]^. Atherosclerosis is the principal pathological basis of major cardiovascular events, including myocardial infarction, ischemic stroke, and peripheral artery disease^[2]^. Its development is driven by the interplay of lipid accumulation, endothelial dysfunction, immune activation, oxidative stress, and vascular remodeling^[2–4]^. Although lipid-lowering and anti-inflammatory therapies have substantially improved cardiovascular outcomes, considerable residual risk persists. Increasing attention to the exposome has highlighted environmental pollutants as potentially important, yet insufficiently characterized, modifiers of atherosclerosis beyond conventional risk factors ^[4,5]^.

Perfluorooctane sulfonate (PFOS) is a prototypical long-chain per- and polyfluoroalkyl substance (PFAS) that has been widely used in industrial and consumer applications, including surfactants, food packaging, textile treatments, electroplating, and firefighting foams^[6,7]^. The exceptional stability of its carbon-fluorine bonds confers environmental persistence, bioaccumulation, and long-range transport. Humans are exposed to PFOS through contaminated food, drinking water, air, and dust, and the compound can persist in the body for prolonged periods^[6–8]^. Epidemiological studies have associated PFAS exposure with dyslipidemia and adverse cardiovascular outcomes, with PFOS particularly linked to elevated cholesterol, adverse lipoprotein profiles, and coronary heart disease risk ^[9–13]^. Experimental studies further suggest that PFOS can aggravate atherosclerotic lesions by promoting macrophage polarization and NF-κB-dependent inflammation^[14]^. However, the systemic mechanisms linking PFOS exposure to atherosclerosis remain incompletely understood.

The gut microbiota is a major regulator of host metabolic and immune homeostasis and represents an important interface between environmental exposures and cardiovascular disease ^[15–17]^. Microbial communities and their metabolites can influence atherosclerosis through multiple pathways, including trimethylamine N-oxide production, short-chain fatty acid metabolism, bile acid biotransformation, epithelial barrier regulation, and endotoxin-driven inflammation ^[15–18]^. Among these mechanisms, bile acids function not only as metabolic substrates but also as signaling molecules that regulate lipid metabolism, intestinal barrier integrity, and immune responses through receptors such as the farnesoid X receptor (FXR) and TGR5 ^[16]^. Consequently, pollutant-induced changes in microbial composition or metabolic capacity may be transmitted to the host through altered bile acid signaling.

PFOS exposure has been shown to perturb gut microbial composition and host metabolic homeostasis ^[19–20]^. More broadly, environmental contaminants can exert selective pressure on intestinal microbial communities and favor bacterial populations capable of tolerating, binding, or metabolizing xenobiotics ^[21–22].^ Nevertheless, it remains unclear whether the gut microbiota is required for PFOS-induced atherosclerosis, which microbial taxa mediate this effect, and how microbial metabolic changes are sensed by the intestinal mucosa to initiate proatherogenic inflammation. In particular, whether bile acid-dependent intestinal FXR signaling can be coupled to innate immune activation in the context of PFOS exposure has not been established.

In this study, we investigated the gut-derived mechanisms underlying PFOS-associated atherosclerosis. Human data showed that fecal PFOS levels were associated with adverse lipid profiles and coronary atherosclerotic burden. Using microbiota depletion and fecal microbiota transplantation, we established a causal contribution of the gut microbiota to the proatherogenic effects of PFOS. We further identified PFOS-enriched *Bacteroides caecimuris* (*B.caecimuris*) as a functional mediator that reduced tauro-ursodeoxycholic acid (TUDCA) levels and aggravated atherosclerosis. Mechanistically, TolC-mediated efflux enhanced the tolerance of *B.caecimuris* to PFOS and supported its intestinal abundance during exposure. The resulting reduction in TUDCA activated intestinal FXR, which directly upregulated *Tlr3* transcription and promoted NF-κB/IRF3 inflammatory signaling, epithelial barrier impairment, and systemic inflammation. Genetic and pharmacological interventions further established TLR3 as a functional downstream effector of intestinal FXR. Together, these findings define a gut microbiota-bile acid-intestinal FXR-TLR3 axis that links PFOS exposure to atherosclerosis progression.

## Results

### PFOS exposure exacerbates atherosclerosis

To evaluate the associations of serum and fecal PFOS with lipid metabolism and coronary atherosclerotic burden, we included 127 participants with available measurements of both serum and fecal PFOS, among whom 82 had coronary angiography-derived Gensini score data (Supplementary Table 1). Partial Spearman correlation analysis, adjusted for age, sex, BMI, smoking status, alcohol consumption, systolic blood pressure, and diastolic blood pressure, showed that serum PFOS was not significantly associated with any lipid-related parameter, including TC, TG, HDL-C, LDL-C, VLDL-C, ApoA1, ApoB, ApoA1/B, Lp(a), and Atherogenic Index (AI). Similarly, no significant association was observed between serum PFOS and the Gensini score(GS score) (*r* = -0.069, *P* = 0.539). In contrast, fecal PFOS was significantly associated with multiple adverse lipid parameters, including TC (*r* = 0.244, *P* = 0.006), TG (*r* = 0.213, *P*= 0.016), LDL-C (*r* = 0.221, *P*= 0.013), VLDL-C (*r* = 0.289, *P* < 0.001), ApoB (*r* = 0.257, *P* = 0.003), ApoA1/B (*r* = -0.255, *P* = 0.004), and AI (*r* = 0.239, *P* = 0.007). Moreover, among the 82 participants with coronary angiography data, fecal PFOS was positively correlated with the Gensini score (*r* = 0.245, *P* = 0.027) (Figure 1A and B).

**Figure 1.**
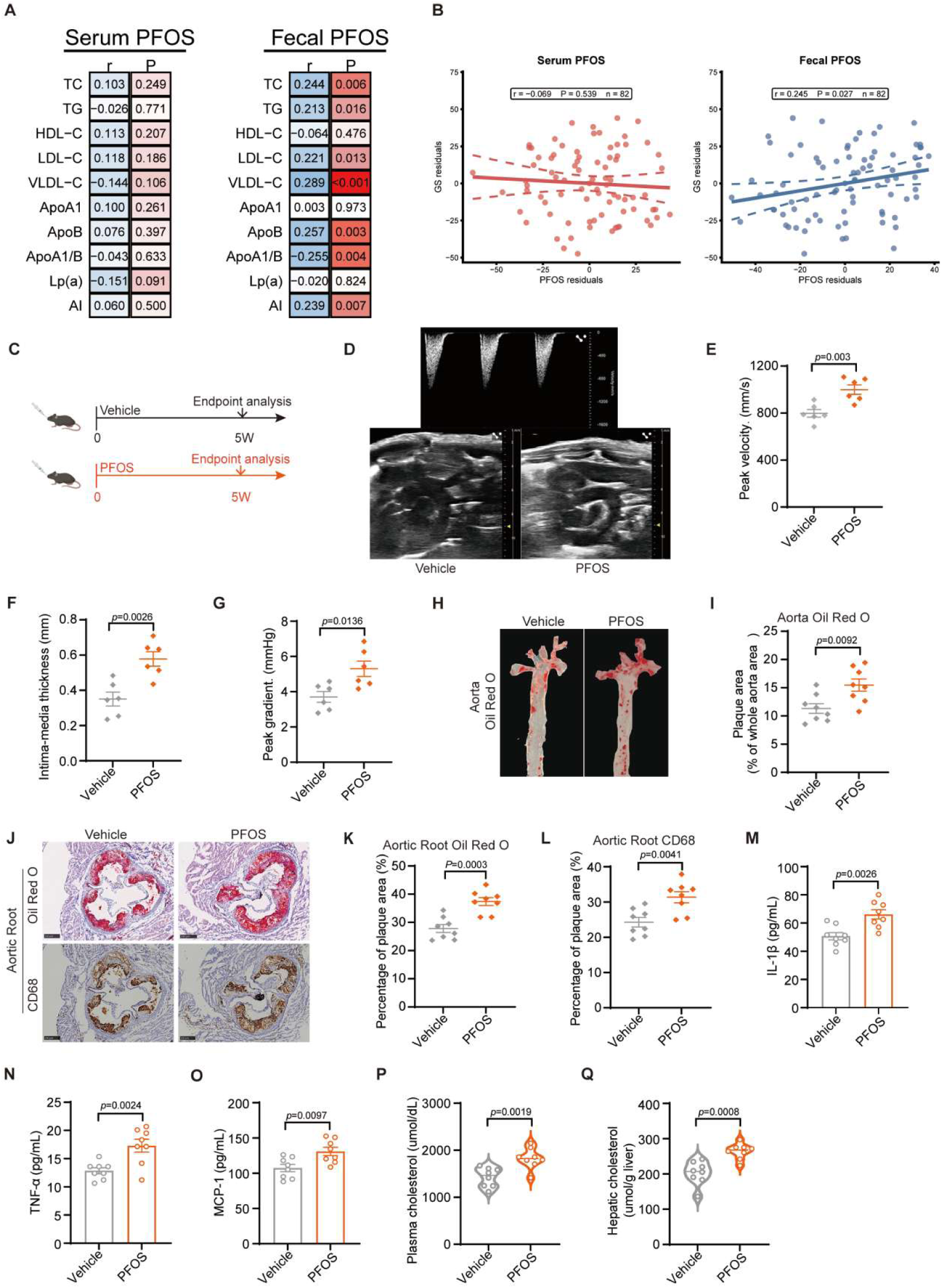
PFOS exposure exacerbates atherosclerosis. **A** and **B**, Associations of human serum and fecal sample PFOS with lipid-related indicators and coronary atherosclerotic burden. The heatmaps on the left show the partial Spearman correlation coefficients (*r*) and *P* values between serum or fecal PFOS levels and lipid-related indicators among participants with paired serum and fecal PFOS measurements (n = 127). The analyzed indicators included TC, TG, HDL-C, LDL-C, VLDL-C, ApoA1, ApoB, ApoA1/B, Lp(a), and AI. The scatter plots on the right show the partial correlations of serum and fecal PFOS with GS score. Points represent covariate-adjusted residuals, solid lines indicate fitted linear trends, and dashed lines indicate the boundaries of the 95% confidence intervals. Correlation analyses were adjusted for age, sex, BMI, smoking status, drinking status, systolic blood pressure, and diastolic blood pressure. The GS score analysis was performed among 82 participants with available coronary angiography data.**C**,Experimental design of chronic PFOS exposure in HCD-fed *ApoE*^−/−^ mice. n = 8 per group. **D-G**, Quantitative analysis of blood flow velocity by ultrasound biomicroscopy (UBM) (top); representative images showing of aorta (bottom) (**D**). Quantitative analysis of the cardiac tract velocity (**E**), quantitative analysis of Intima-media thickness (**F**) and quantitative analysis of the aortic peak grad (**G**). n = 6 per group. **H** and **I**, Representative images of aortas stained with Oil Red O (**H**) and quantification of the lesions (**I**). n = 8 per group. **J**, Representative images of Oil Red O staining and CD68^+^ macrophage staining of aortic root cross-sections. Scale bar, 250 μm. n = 8 per group. **K** and **L**,Quantification analysis of Oil Red O staining (**J**) and CD68^+^ macrophage staining (**K**) of aortic roots. n = 8 per group. **M-O**, Levels of the proinflammatory cytokines IL-1β (**M**), TNF-α (**N**), and MCP-1 (**O**) in the plasma. n = 8 per group. **P** and **Q**, Plasma (**P**) and Hepatic (**Q**) cholesterol levels of the indicated mice. n = 8 per group. Data are presented as mean ± SEM. Each dot represents one biological replicate. *P* values are indicated in the panels. Two-tailed Student’s t test (**E-G**,**I**,**K-Q**). Source data are provided as a Source Data file.

**Supplementary Figure 1.**
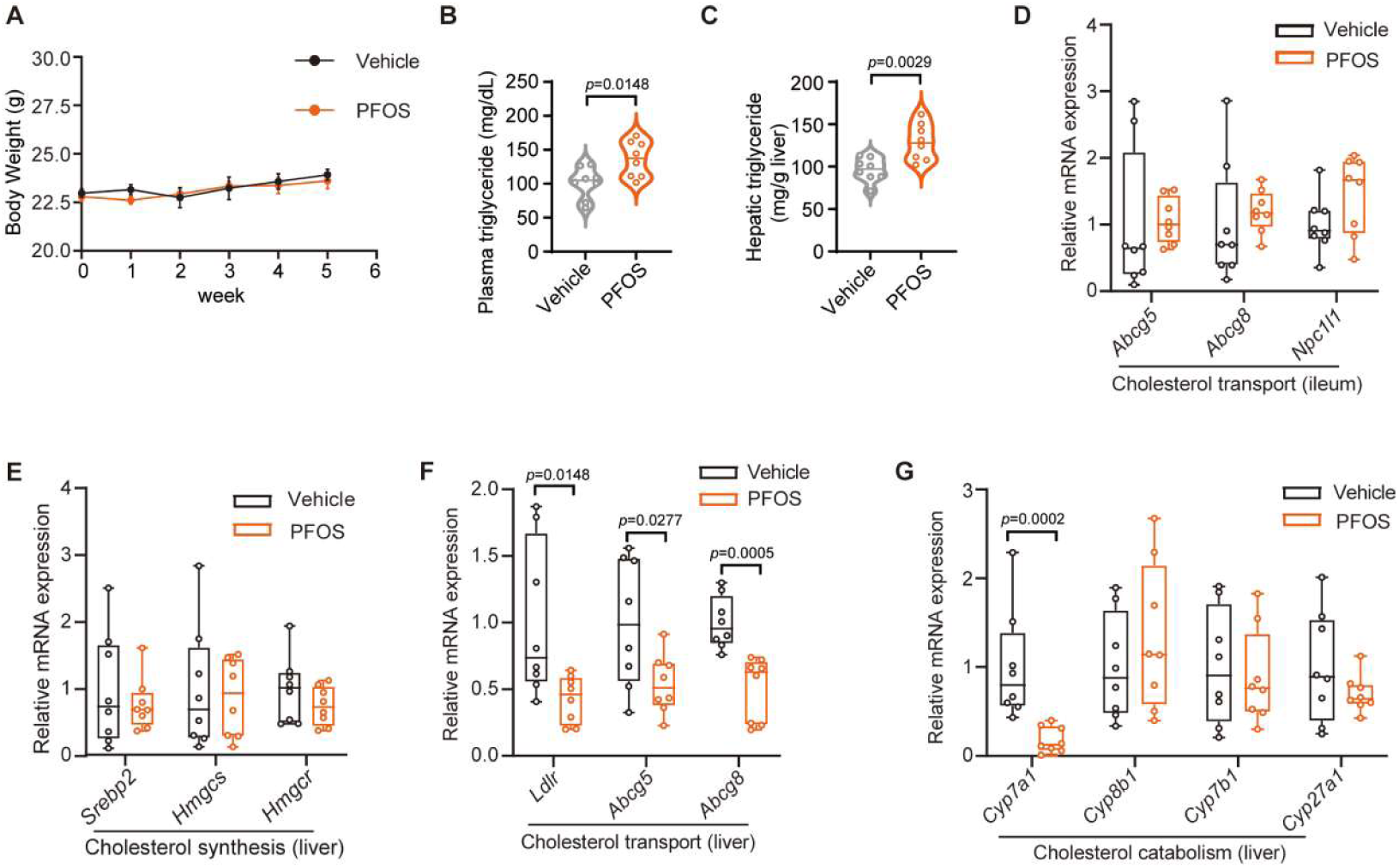
PFOS exposure exacerbates atherosclerosis. Related to Figure 1. **A**, Body weights.n = 8 per group. **B** and **C**, Plasma (**B**) and hepatic (**C**) triglyceride levels of the indicated mice.n = 8 per group. **D**, Levels of mRNAs encoded by genes related to cholesterol transport in the ileum.n = 8 per group. **E**-**G**, Levels of mRNAs encoded by genes related to cholesterol synthesis (**E**),cholesterol transport (**F**),and cholesterol elimination (**G**) in the liver.n = 8 per group.Data are presented as mean ± SEM. Each dot represents one biological replicate. *P* values are indicated in the panels. Two-tailed Student’s t test (**B-C**,**F**).Mann-Whitney U test (**D-G**).Source data are provided as a Source Data file.

To further investigate the effect of PFOS on the development of atherosclerosis, *ApoE*^−/−^ mice were randomly assigned to two groups and administered PFOS or vehicle by oral gavage for 5 weeks (Figure 1C). To minimize the confounding effect of obesity, we used a high-cholesterol diet (HCD) regimen that did not induce significant body weight gain (Supplementary Figure 1A). At the end of the intervention, cardiac blood flow, peak pressure gradient, and aortic intima-media thickness (IMT) were assessed by color Doppler ultrasound microscopy. Compared with vehicle-treated controls, PFOS-treated mice exhibited significantly worsened hemodynamic parameters and increased aortic IMT (Figure 1D-G). Oil Red O staining further revealed that PFOS-treated mice fed an HCD developed more severe atherosclerotic lesions throughout the entire aorta and in aortic root sections than control mice (Figure 1H-K). Given that macrophages represent a predominant immune cell population within atherosclerotic plaques and play a critical role in lesion initiation and progression, we next performed immunostaining for the macrophage marker CD68 in aortic root plaques. The proportion of CD68-positive area within plaques was significantly increased in PFOS-treated mice, indicating enhanced macrophage infiltration in the lesion area (Figure 1J and L). Consistently, circulating levels of inflammatory cytokines were also markedly elevated in the plasma of PFOS-treated mice compared with controls (Figure 1M-O).

In addition, PFOS-treated mice fed an HCD showed higher levels of cholesterol and triglycerides in both plasma and liver than vehicle-treated controls (Figure 1P,Q and Supplementary Figure 1B,C). Correspondingly, no significant differences were observed in the mRNA expression of genes involved in intestinal cholesterol transport or hepatic cholesterol synthesis between the two groups. However, hepatic expression of the cholesterol transport-related genes *Ldlr, Abcg5, and Abcg8* was modestly reduced in PFOS-treated mice (Supplementary Figure 1D-F). Moreover, expression of *Cyp7a1*, a key enzyme involved in cholesterol catabolism, was also suppressed in the liver of PFOS-treated mice (Supplementary Figure 1G). Together, these data indicate that PFOS exacerbates HCD-driven atherosclerosis in *ApoE*^−/−^ mice, accompanied by enhanced vascular inflammation, macrophage accumulation, systemic dyslipidemia, and altered hepatic cholesterol handling.

### The gut microbiota mediates PFOS-induced exacerbation of atherosclerosis

To determine whether the gut microbiota plays a critical role in PFOS-induced atherosclerosis progression, we first depleted the gut microbiota in *ApoE*^−/−^ mice using broad-spectrum antibiotics (Supplementary Figure 2A). After antibiotic treatment, Oil Red O staining of the whole aorta and aortic root sections showed that PFOS exposure no longer exacerbated atherosclerotic lesion formation. Consistently, macrophage infiltration within aortic root plaques was not significantly different between the two groups (Figure 2A-E). After antibiotic-mediated microbiota depletion, circulating inflammatory cytokine levels did not differ significantly between PFOS-treated and vehicle-treated mice (Figure 2F-H). Notably, antibiotic treatment abolished the differences in plasma and hepatic cholesterol and triglyceride levels between PFOS-exposed and control mice (Figure 2I,J and Supplementary Figure 2B,C).

**Figure 2.**
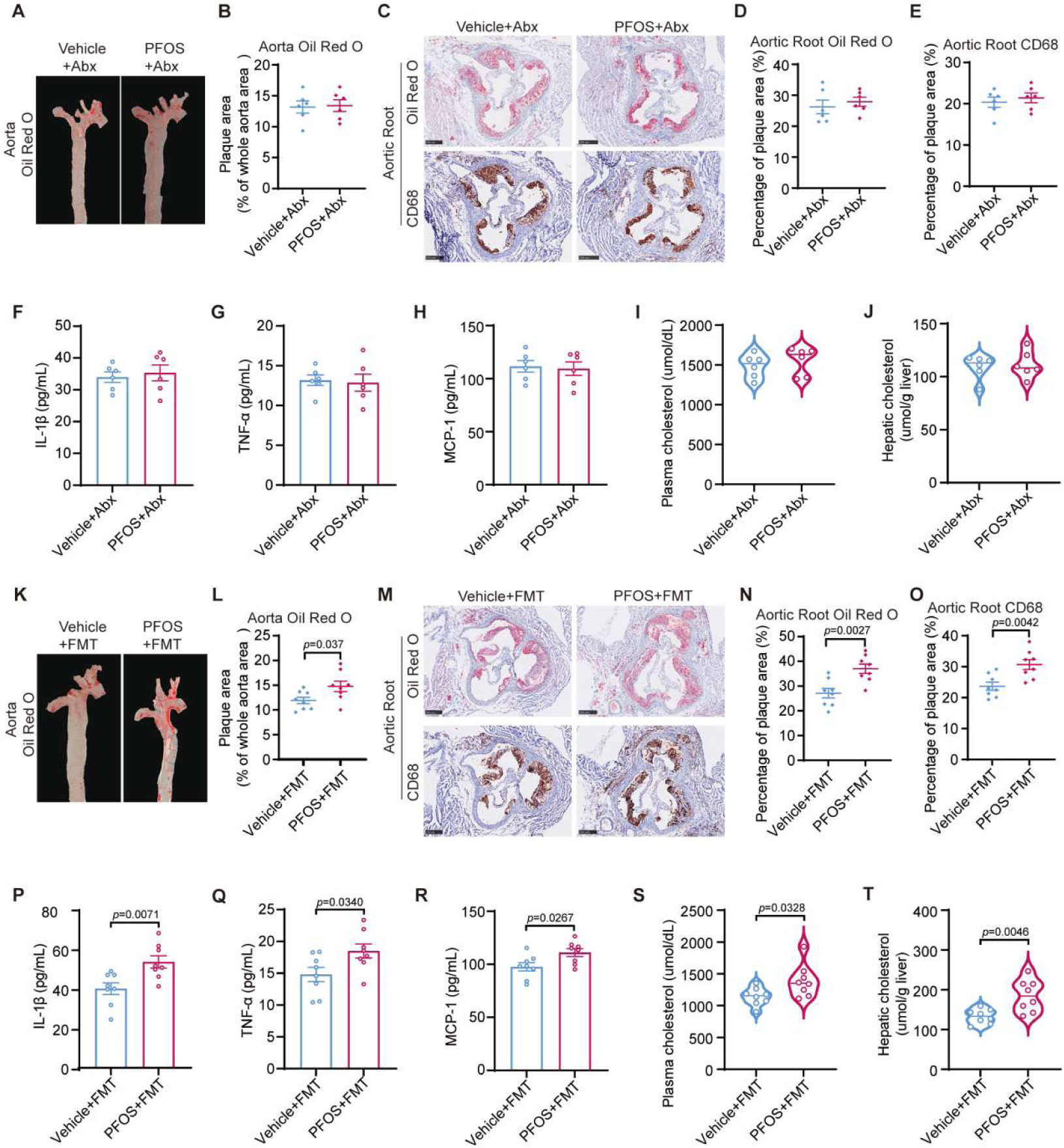
The gut microbiota mediates PFOS-induced exacerbation of atherosclerosis. **A** and **B**, Representative images of aortas stained with Oil Red O (**A**) and quantification of the lesions (**B**). n = 6 per group. **C**,Representative images of Oil Red O staining and CD68^+^ macrophage staining of aortic root cross-sections. Scale bar, 250μm. n = 6 per group. **D** and **E**, Quantification analysis of Oli Red O staining (**D**) and CD68^+^ macrophage staining (**E**) of aortic roots.n = 6 per group. **F-H**, Levels of the proinflammatory cytokines IL-1β (**F**), TNF-α (**G**), and MCP-1 (**H**) in the plasma.n = 6 per group. **I** and **J**, Plasma (**I**) and hepatic (**J**) cholesterol levels of the indicated mice.n = 6 per group. **K** and **L**, Representative images of aortas stained with Oil Red O (**K**) and quantification of the lesions (**L**). n = 8 per group. **M**,Representative images of Oil Red O staining and CD68^+^ macrophage staining of aortic root cross-sections. Scale bar, 250 μm. n = 8 per group. **N** and **O**, Quantification analysis of Oil Red O staining (**N**) and CD68^+^ macrophage staining (**O**) of aortic roots. n = 8 per group. **P-R**, Levels of the proinflammatory cytokines IL-1β (**P**), TNF-α (**Q**), and MCP-1 (**R**) in the plasma.n = 8 per group. **S** and **T**, Plasma (**S**) and hepatic (**T**) cholesterol levels of the indicated mice. n = 8 per group. Data are presented as mean±SEM. Each dot represents one biological replicate. *P* values are indicated in the panels. Two-tailed Student’s t test (**B**,**D-J**,**L**,**N-T**). Source data are provided as a Source Data file.

**Supplementary Figure 2.**
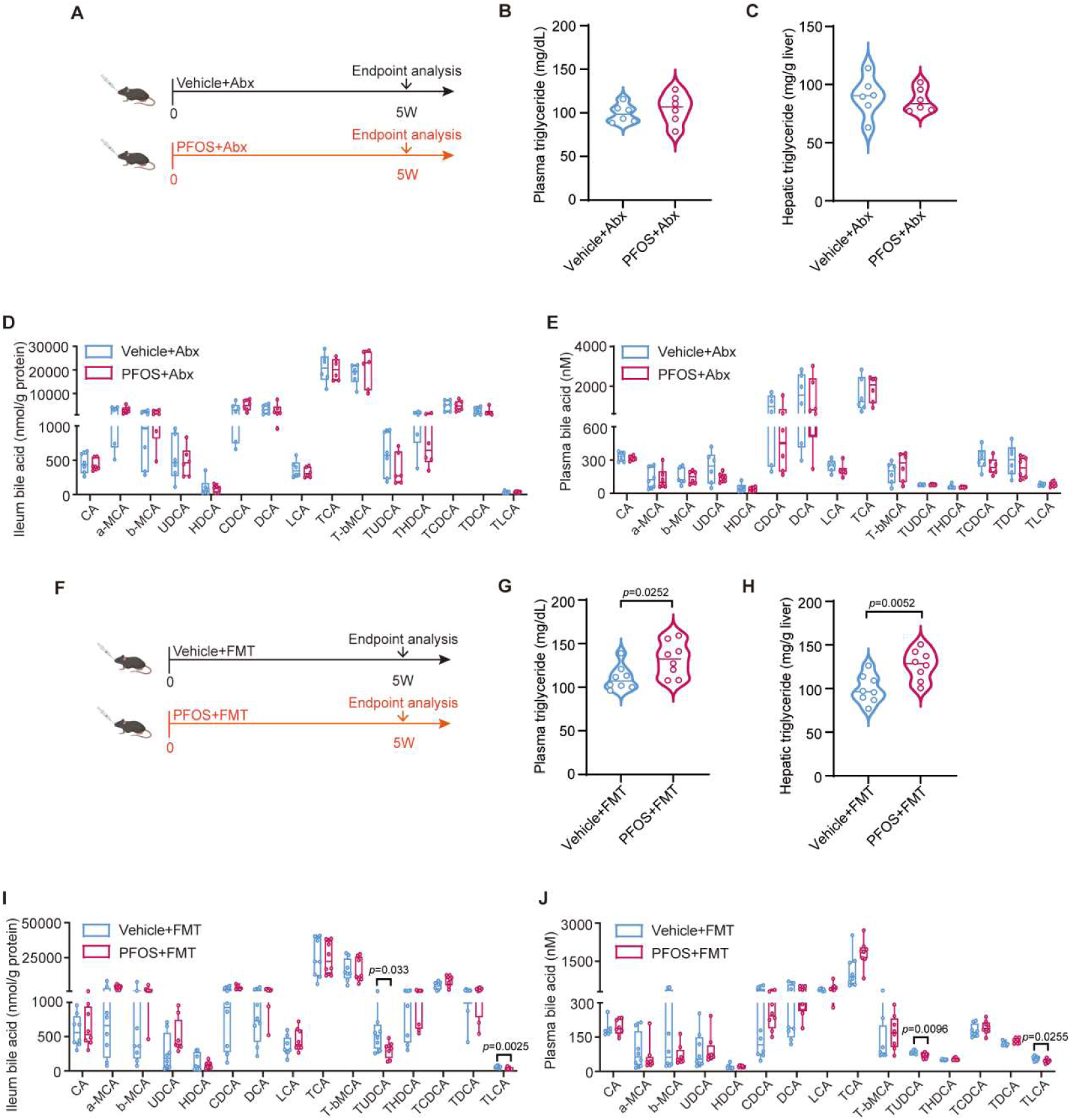
The gut microbiota mediates PFOS-induced exacerbation of atherosclerosis. Related to Figure 2. **A**, Schematic of the Abx experimental procedure. n = 6 per group. **B** and **C**, Plasma (**B**) and hepatic (**C**) triglyceride levels of the indicated mice.n = 6 per group. **D** and **E**, Bile acid concentrations in the ileum (**D**) and plasma (**E**) in vehicle + Abx and PFOS + Abx mice fed a HCD.n = 6 per group. **F**, Schematic of the FMT experimental model.n = 8 per group. **G** and **H**, Plasma (**G**) and hepatic (**H**) triglyceride levels of the indicated mice.n = 8 per group. **I** and **J**, Bile acid concentrations in the ileum (**I**) and plasma (**J**) in vehicle + FMT and PFOS + FMT mice fed a HCD.n = 8 per group.Data are presented as mean ± SEM. Each dot represents one biological replicate. *P* values are indicated in the panels. Two-tailed Student’s t test(**B-E**,**G-J**).Source data are provided as a Source Data file.

To further determine whether the PFOS-induced atherogenic phenotype could be transferred through the gut microbiota, we performed fecal microbiota transplantation (FMT) using fecal microbiota from PFOS-or vehicle-treated donor mice (Supplementary Figure 2F). Recipient mice colonized with microbiota from PFOS-treated donors exhibited significantly larger atherosclerotic lesion areas in both the whole aorta and aortic root sections compared with those receiving microbiota from vehicle-treated donors (Figure 2K-N). Moreover, macrophage accumulation in aortic root plaques was markedly increased in the PFOS-FMT group (Figure 2M,O). Consistently, circulating inflammatory cytokine levels were significantly elevated in mice receiving PFOS-associated microbiota (Figure 2P-R). In parallel, plasma cholesterol and triglyceride levels were also higher in the PFOS-FMT group than in the Vehicle-FMT group (Figure 2S,T and Supplementary Figure 2G and H).

Together, these findings demonstrate that antibiotic-mediated microbiota depletion abrogates PFOS-induced aggravation of atherosclerosis, whereas transplantation of PFOS-associated microbiota is sufficient to recapitulate this phenotype. These results identify the gut microbiota as a critical mediator of PFOS-driven metabolic disturbance and atherosclerosis progression.

### PFOS-enriched *B.caecimuris* alters TUDCA homeostasis and exacerbates atherosclerosis

To characterize PFOS-induced alterations in the gut microbiota, cecal contents from PFOS-exposed and vehicle-treated mice were subjected to shotgun metagenomic sequencing. PFOS exposure altered α-diversity, and principal coordinate analysis further revealed a clear separation between the two groups, indicating a marked shift in overall gut microbial community structure (Figure 3A,B). At the order level, linear discriminant analysis effect size (LEfSe) identified *Bacteroidales* as enriched in PFOS-exposed mice and as having the largest LDA effect size among the bacterial orders that differed between PFOS-exposed and vehicle-treated mice (LDA score = 5.02; Figure 3C and D). Because the LDA score reflects the discriminatory effect size of a taxon rather than its relative abundance, we next examined order-level microbial composition. *Bacteroidales* represented a major component of the gut microbiota and exhibited a higher relative abundance in PFOS-exposed mice than in vehicle-treated controls (Figure 3D). Together, these complementary analyses demonstrated that PFOS exposure markedly increased the representation of *Bacteroidales* in the gut microbiota.

**Figure 3.**
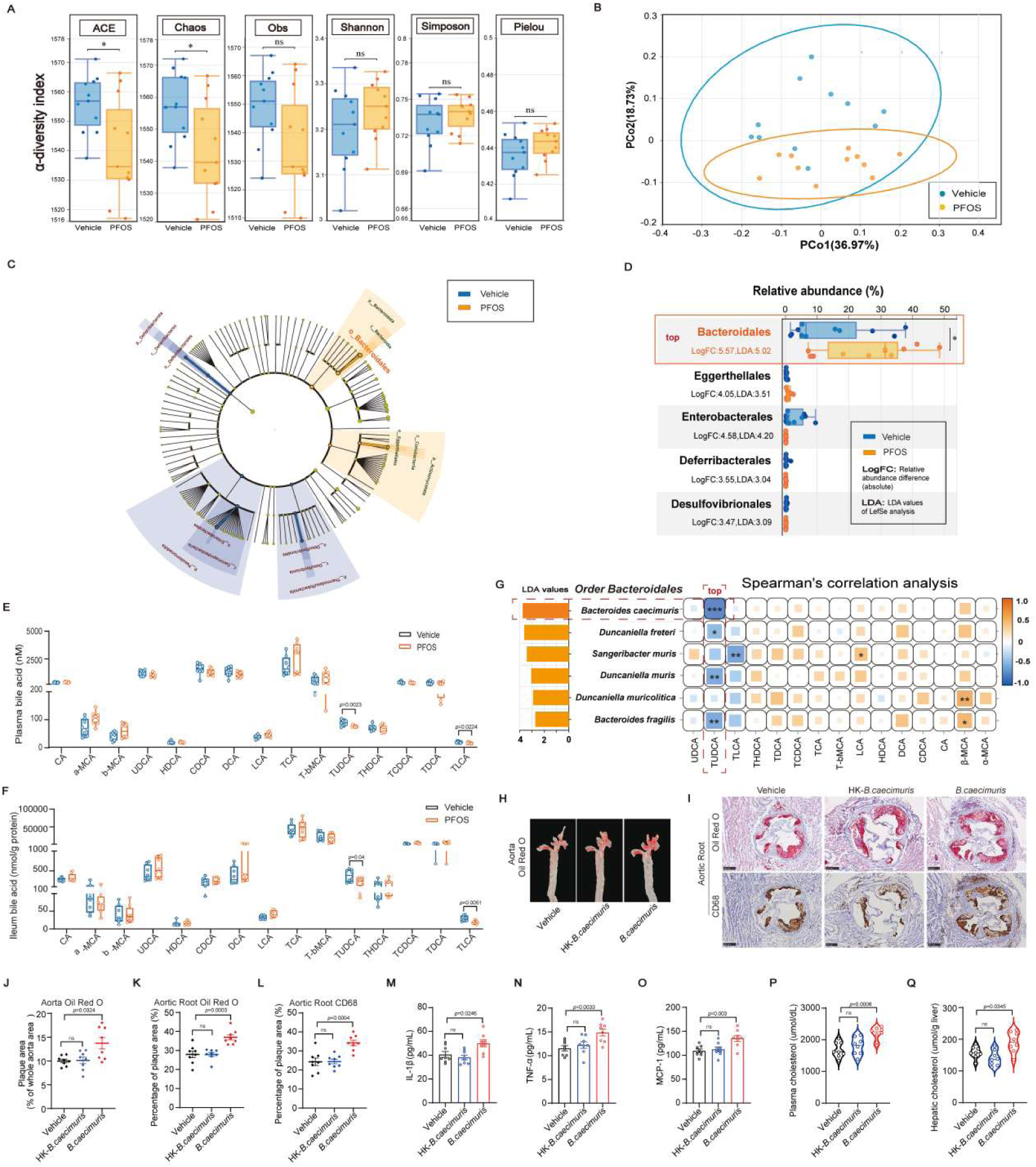
PFOS-enriched B.caecimuris alters TUDCA homeostasis and exacerbates atherosclerosis. **A** and **B**, α-diversity of the gut microbiota between the vehicle and PFOS groups, as indicated by the obs, ACE, Chaos, and Shannon indices. Principal coordinate analysis (PCoA) illustrates the beta diversities (Bray-Curtis distances) of microbial taxonomy species. n = 11 per group. **C**, Taxonomic cladogram generated from linear discriminant analysis (LDA) effect size (LEfSe) of metagenomic sequencing data. Blue indicates enriched taxa in the vehicle group. Yellow indicates enriched taxa in the PFOS exposure group. **D**, Abundances (percent) of microbial species between the two groups based on metagenomics data. Each dot represents an individual mouse. N = 11 per group. **E** and **F**, Bile acid concentrations in the plasma (**E**) and ileum (**F**) in vehicle and PFOS mice fed a HCD. n = 8 per group. **G**, The association between changes in microbial species (with LDA values > 3) belonging to order Bacteroidetes and host bile acid levels. Bar graph on the left denotes the LDA values of each species. Yellow indicates enriched taxa in the PFOS exposure group. The right heatmap denotes the Spearman’s coefficient of bile acid levels with the abundance of microbial species. **H** and **J**, Representative images of aortas stained with Oil Red O (**H**) and quantification of the lesions (**J**). n = 8 per group. **I**, Representative images of Oil Red O staining and CD68^+^ macrophage staining of aortic root cross-sections. Scale bar, 250μm. n = 8 per group. **K** and **L**, Quantification analysis of Oil Red O staining (**K**) and CD68^+^ macrophage staining (**L**) of aortic roots. n = 8 per group. **M-O**, Levels of the proinflammatory cytokines IL-1β (**M**), TNF-α (**N**), and MCP-1 (**O**) in the plasma. n = 8 per group. **P** and **Q**, Plasma (**P**) and hepatic (**Q**) cholesterol levels of the indicated mice. n = 8 per group. Data are presented as mean±SEM. Each dot represents one biological replicate. *P* values are indicated in the panels. In the box plots, the line indicates the median; the box represents the interquartile range (IQR, middle 50% of the data); whiskers extend to data points within 1.5×IQR; points beyond are outliers (**A-G**). Two-sided Mann-Whitney U test (**A** and **D**). Two-tailed Student’s t test (**E**-**F**) Spearman’s rank tests (**G**). Kruskal-Wallis test with Dunn’s post hoc test (**J**). One-way ANOVA with Tukey’s post hoc test (**K-Q**). Source data are provided as a Source Data file.

**Supplementary Figure 3.**
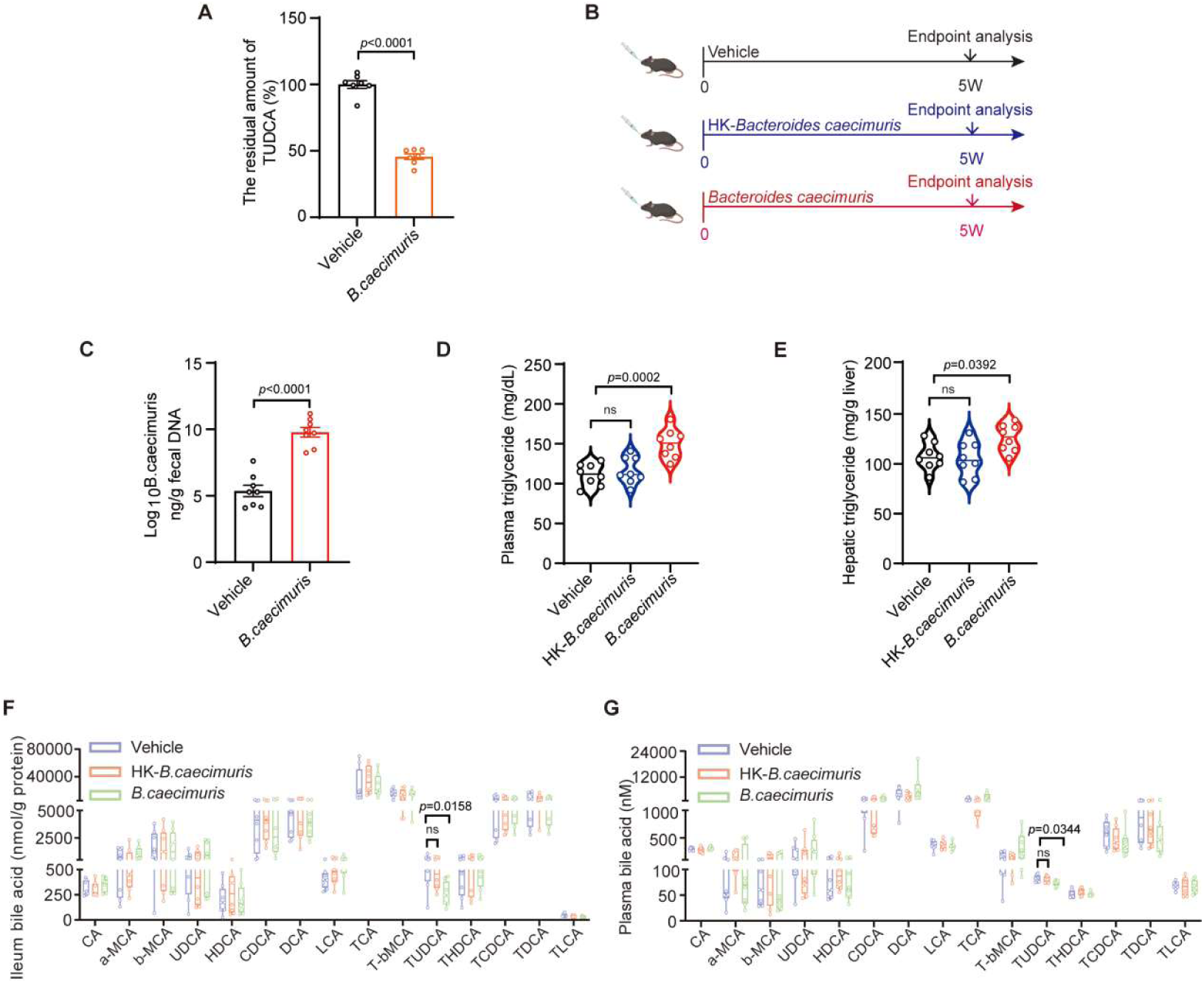
PFOS-enriched B.caecimuris alters TUDCA homeostasis and exacerbates atherosclerosis. Related to Figure 3. **A**, The hydrolysis efficiency of TUDCA by *B.caecimuris*(7 biological replicates for each group). **B**, Experimental design of chronic B.caecimuris gavage in HCD-fed *ApoE*^−/−^ mice.n = 8 per group. **C**, Changes in qPCR-based absolute abundances of fecal *B.caecimuris* in mouse.n = 8 per group. **D** and **E**, Plasma (**D**) and hepatic (**E**) triglyceride levels of the indicated mice.n = 8 per group. **F** and **G**, Bile acid concentrations in the ileum (**F**) and plasma (**G**) in vehicle, HK-*B.caecimuris* and *B.caecimuris* fed a HCD. n = 8 per group.Data are presented as mean ± SEM. Each dot represents one biological replicate. *P* values are indicated in the panels. Two-tailed Student’s t test(**A** and **C**).One-way ANOVA with Tukey’s post hoc test(**D** and **E**). Kruskal-Wallis test with Dunn’s post hoc test (**F** and **G**).Source data are provided as a Source Data file.

Members of *Bacteroidales* are known to harbor bile salt hydrolases (BSHs), which catalyze the deconjugation of conjugated bile acids into unconjugated bile acids and their corresponding amino acid moieties, thereby contributing to remodeling of the intestinal bile acid pool ^[23–25]^. In addition, disruption of bile acid metabolism and signaling has been closely associated with cardiometabolic diseases ^[26–27]^. We therefore investigated whether PFOS-induced microbial remodeling was accompanied by alterations in bile acid homeostasis. Targeted bile acid profiling showed that PFOS exposure significantly reduced the concentrations of two taurine-conjugated bile acids, tauroursodeoxycholic acid (TUDCA) and taurolithocholic acid (TLCA), in both plasma and ileal (Figure 3E and F).

We next performed Spearman correlation analysis between ileal bile acid levels and PFOS-enriched *Bacteroidales* species with LDA scores greater than 3. Multiple species-bile acid associations were observed. Notably, *B.caecimuris* exhibited the largest LDA effect size among the examined *Bacteroidales* species and was significantly inversely correlated with TUDCA levels in the ileal (Figure 3G). Together with the consistent reduction in TUDCA following PFOS exposure, these findings prioritized a candidate *B.caecimuris*-TUDCA link for subsequent functional investigation.

To determine whether *B.caecimuris* could directly contribute to TUDCA depletion, we assessed its capacity to metabolize TUDCA in vitro. Incubation with viable *B.caecimuris* resulted in substantial hydrolysis and depletion of TUDCA, demonstrating a direct metabolic interaction between *B.caecimuris* and TUDCA (Supplementary Figure 3A). Together with the inverse association between *B.caecimuris* abundance and ileal TUDCA levels, this finding provided a rationale for evaluating both the pathogenic potential of *B.caecimuris* and its effect on TUDCA homeostasis in vivo.

Having established that microbiota from PFOS-exposed donors was sufficient to aggravate atherosclerosis in recipient mice, we next asked whether the PFOS-enriched species *B.caecimuris* could independently reproduce this phenotype in the absence of direct PFOS exposure. HCD-fed *ApoE*^⁻/⁻^ mice were orally administered viable *B. caecimuris*, heat-killed *B.caecimuris* (HK-*B.caecimuris*), or vehicle for 5 weeks (Supplementary Figure 3B). At the experimental endpoint, qPCR analysis confirmed successful intestinal colonization, as reflected by a marked increase in the fecal abundance of *B.caecimuris* in mice receiving the viable bacterium (Supplementary Figure 3C).

Administration of viable *B.caecimuris* significantly increased atherosclerotic lesion burden in both the whole aorta and aortic root compared with vehicle treatment (Figure 3H-K). Consistently, CD68 immunostaining revealed increased macrophage accumulation within aortic root lesions, and circulating pro-inflammatory cytokine levels were also elevated in mice administered viable *B.caecimuris* (Figure 3I,L–O). In parallel, viable *B.caecimuris* increased plasma and hepatic cholesterol levels, together with elevated plasma and hepatic triglyceride levels (Figure 3P,Q and Supplementary Figure 3D and E). In contrast, HK-*B.caecimuris* did not significantly alter these atherosclerotic, inflammatory, or metabolic parameters relative to vehicle controls. These findings demonstrate that viable *B.caecimuris* is sufficient to exacerbate HCD-induced atherosclerosis and associated systemic inflammatory and metabolic abnormalities independently of direct PFOS exposure, indicating that its proatherogenic effect requires bacterial viability.

We next determined whether administration of *B.caecimuris* reproduced the TUDCA alteration observed following PFOS exposure. Compared with vehicle-treated mice, mice receiving viable *B.caecimuris* exhibited significantly reduced TUDCA levels in both ileal and plasma, whereas HK-*B.caecimuris* did not significantly alter TUDCA levels relative to vehicle controls (Supplementary Figure 3F and G). Notably, no other measured bile acids showed consistent changes in both compartments. Together, these findings demonstrate that viable *B.caecimuris* is sufficient to exacerbate HCD-induced atherosclerosis independently of direct PFOS exposure. In parallel, viable *B.caecimuris* consistently reduced TUDCA levels in both ileal and plasma, further supporting its role in the bile acid remodeling observed following PFOS exposure.

### TolC-mediated efflux confers PFOS tolerance and supports intestinal colonization of *B.caecimuris*

Having established that *B.caecimuris* was enriched following PFOS exposure and was sufficient to aggravate atherosclerosis, we next investigated the mechanism underlying its increased abundance. We hypothesized that *B.caecimuris* may possess an intrinsic capacity to tolerate PFOS, thereby facilitating its maintenance in the gut during PFOS exposure. To test this possibility, we determined the minimum inhibitory concentration (MIC) of PFOS against *B.caecimuris*. The MIC was 256 μg/mL, indicating that *B.caecimuris* can withstand relatively high concentrations of PFOS (Figure 4A). This finding raised the possibility that PFOS tolerance contributes to its increased intestinal abundance under PFOS exposure.

**Figure 4.**
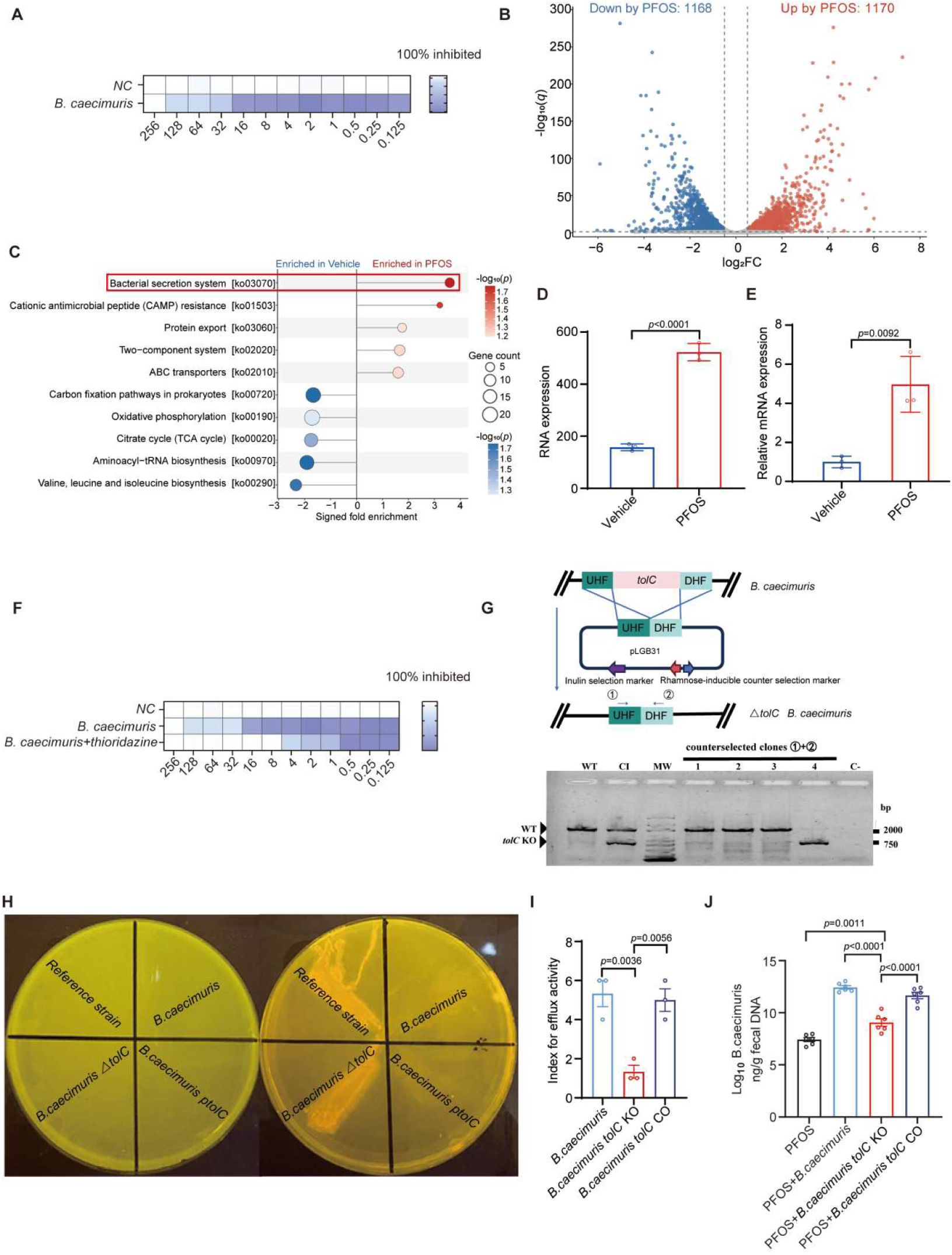
TolC-mediated efflux confers PFOS tolerance and supports intestinal colonization of B.caecimuris. **A**, Minimum inhibitory concentration of *B.caecimuris* towards PFOS. **B**, Volcano plot of differentially expressed genes between PFOS and Vehicle groups. Differential genes were defined as q < 0.05 and |log₂FC| ≥ 0.5. Red and blue indicate genes upregulated and downregulated by PFOS, respectively; gray indicates non-significant genes. **C**, Bidirectional bubble plot of representative KEGG pathways enriched among PFOS-upregulated or PFOS-downregulated genes. Positive and negative signed fold-enrichment values indicate enrichment in PFOS and Vehicle groups, respectively. Bubble size represents gene count, and color intensity represents − log_10_(P). **D**, RNA expression of TolC family protein gene of *B.caecimuris* exposed to PFOS. **E**, qPCR of TolC family protein gene of *B.caecimuris* exposed to PFOS. **F**, Minimum inhibitory concentration of *B.caecimuris* towards PFOS with and without thioridazine. **G**, Diagram of TolC family protein gene knock out. **H**, Ethidium Bromide-agar Cartwheel test of wild type, TolC family protein gene knock out and complemented *B.caecimuris.* **I**, Index for efflux activity determined by MC_EtBr_ (target strain)-MC_EtBr_ (REF)/MC_EtBr_ (REF). **J,** Changes in qPCR-based absolute abundances of fecal *B.caecimuris* in mouse. n = 6 per group. Data are presented as mean ± SEM. Each dot represents one biological replicate. *P* values are indicated in the panels. Two-tailed Student’s t test (**D** and **E**). One-way ANOVA with Tukey’s post hoc test (**I** and **J**). Source data are provided as a Source Data file.

**Supplementary Figure 4.**
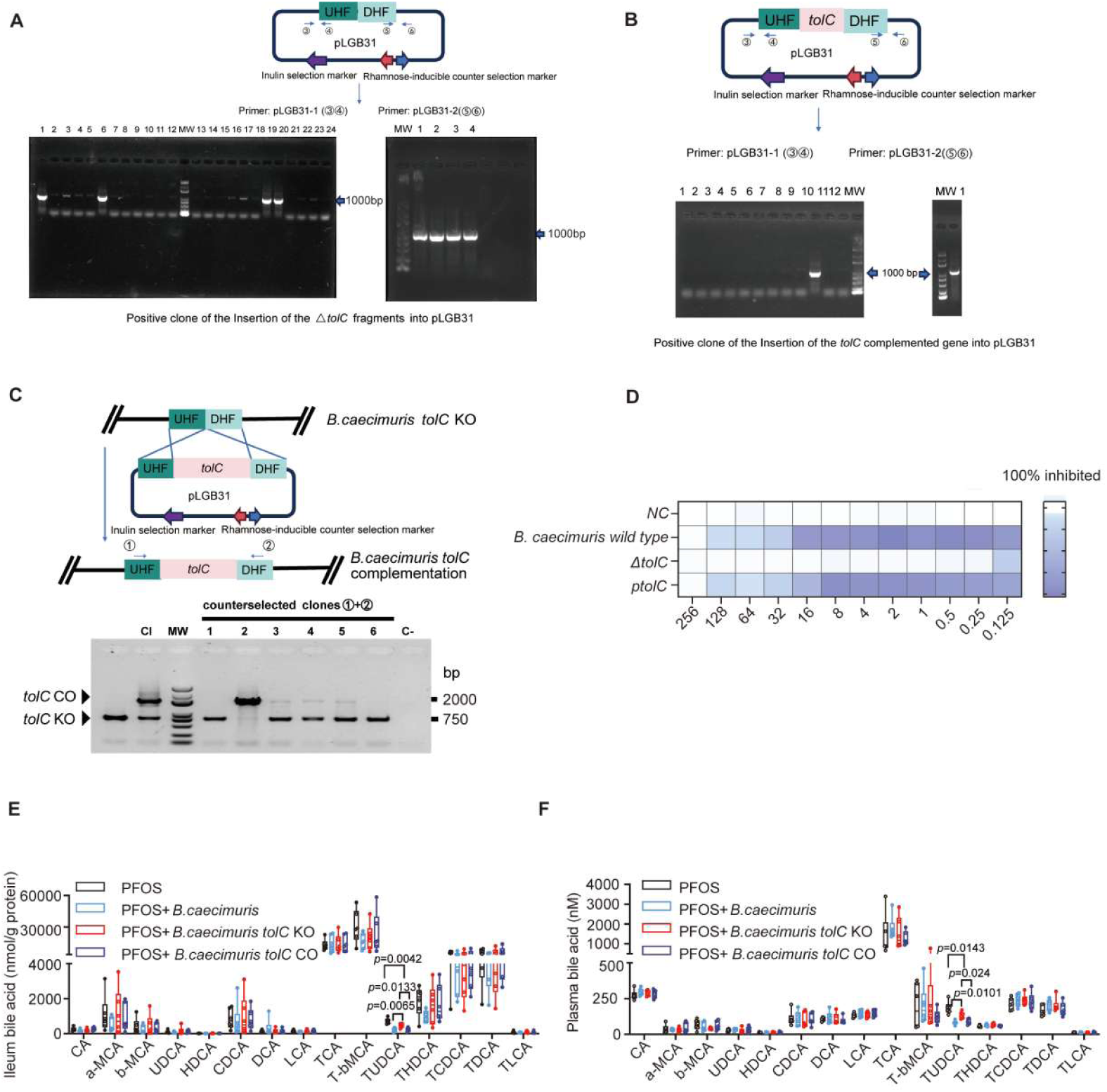
TolC-mediated efflux confers PFOS tolerance and supports intestinal colonization of B. caecimuris. Related to Figure 4. **A**, Selection of △*tolC* positive pLGB31 clones. **B**, Selection of *tolC* complementation pLGB31 clones. **C**, Selection of *tolC* complemented *B.caecimuris.* **D**, Minimum inhibitory concentration of wild type, △*tolC* and *tolC* complemented *B. caecimuris*. **E** and **F**, Bile acid concentrations in the ileum (**E**) and plasma (**F**).n = 6 per group.The median with first and third quartiles in box and whiskers. One-way ANOVA with Tukey’s post hoc test (**E** and **F**).Source data are provided as a Source Data file.

To identify bacterial responses potentially underlying PFOS tolerance, we performed RNA sequencing of *B.caecimuris* following PFOS exposure. Using thresholds of *q* < 0.05 and |log₂FC| ≥ 0.5, differential expression analysis identified 1,170 upregulated and 1,168 downregulated genes in PFOS-exposed bacteria compared with vehicle-treated controls (Figure 4B). KEGG enrichment analysis revealed distinct functional profiles among the upregulated and downregulated gene sets (Figure 4C). The bacterial secretion system was the most significantly enriched pathway among the PFOS-upregulated genes, whereas genes involved in several central metabolic processes were predominantly enriched among the downregulated genes. These findings suggested that enhanced secretion or export capacity may contribute to the adaptation of *B.caecimuris* to PFOS exposure.

We therefore examined genes associated with the bacterial secretion system. Among the PFOS-upregulated genes, the TolC family protein-encoding gene *A4V03_RS15910* (designated *tolC*) was markedly induced and was prioritized for further investigation because of the established role of TolC family proteins in bacterial efflux. The PFOS-induced upregulation of *tolC* observed in the RNA-seq dataset was independently confirmed by qPCR (Figure 4D and E), supporting a potential role for TolC-mediated efflux in the adaptation of *B.caecimuris* to PFOS exposure.

To determine whether bacterial efflux functionally contributes to PFOS tolerance, we pharmacologically inhibited efflux activity using thioridazine. Thioridazine treatment reduced the MIC of PFOS against *B.caecimuris* by 32-fold, indicating that bacterial efflux contributes substantially to PFOS tolerance (Figure 4F). Because pharmacological inhibition does not identify the specific efflux component responsible for this effect, we next directly examined the role of *tolC* using genetic approaches.

A *tolC*-knock out strain (Δ*tolC*) and a *tolC*-complemented strain were generated and experimentally validated (Figure 4G and Supplementary Figure 4A-C). Efflux activity was subsequently evaluated using an ethidium bromide (EtBr) agar cartwheel assay. The Δ*tolC* strain showed greater EtBr retention than the wild-type and *tolC*-complemented strains, consistent with impaired efflux activity following *tolC* deletion (Figure 4H and I). In parallel, the MIC of PFOS against the Δ*tolC* strain was lower than those observed for the wild-type and *tolC*-complemented strains (Supplementary Figure 4D). Together, these pharmacological and genetic findings support an important role for TolC-mediated efflux in the tolerance of *B.caecimuris* to PFOS.

We next investigated whether TolC contributes to the intestinal abundance of *B.caecimuris* under PFOS exposure in vivo. HCD-fed *ApoE*^⁻/⁻^ mice exposed to PFOS were orally administered wild-type, Δ*tolC*, or *tolC*-complemented *B.caecimuris*, with a bacteria-free PFOS-exposed group serving as the control. During ongoing bacterial administration, qPCR analysis showed that the fecal absolute abundance of the Δ*tolC* strain was significantly lower than that of both the wild-type and *tolC*-complemented strains (Figure 4J). These findings indicate that TolC-mediated PFOS tolerance supports the intestinal colonization and maintenance of *B.caecimuris* abundance during PFOS exposure.

Finally, we assessed whether the reduced abundance of the Δ*tolC* strain was accompanied by an attenuated effect on bile acid homeostasis. Administration of wild-type and *tolC*-complemented *B.caecimuris* reduced TUDCA levels in both ileal and plasma, whereas the Δ*tolC* strain showed a diminished capacity to reduce TUDCA (Supplementary Figure 4E and F). Other measured bile acids were not significantly altered. Because PFOS exposure under *tolC*-deficient conditions reduced the intestinal abundance of *B.caecimuris*, the attenuated TUDCA response most likely reflects this reduced bacterial abundance rather than a direct regulation of TUDCA metabolism by TolC itself. Collectively, these results indicate that TolC-mediated efflux enhances PFOS tolerance, thereby supporting the intestinal abundance of *B.caecimuris* and its associated effect on TUDCA homeostasis under PFOS exposure.

### *Tlr3* is a direct transcriptional target of intestinal FXR

Our preceding analyses showed that PFOS exposure reshaped the gut microbiota and altered bile acid homeostasis in association with aggravated atherosclerosis. To characterize the intestinal response to these changes, we performed RNA sequencing of ileal tissues from HCD-fed *ApoE*^⁻/⁻^ mice exposed to PFOS or vehicle. Principal component analysis revealed clear separation between the two groups, which was confirmed by PERMANOVA (R² = 0.312, *P* = 0.0075; Supplementary Figure 5A). Differential expression analysis identified 276 upregulated and 188 downregulated genes in PFOS-exposed mice using thresholds of |log₂FC| > 1 and *P* < 0.05 (Supplementary Figure 5B). Gene set enrichment analysis showed concurrent enrichment of pathways related to bile acid signaling and transport, lipid metabolism, acute inflammatory responses, macrophage cytokine production, and endolysosomal TLR signaling (Figure 5A), indicating coordinated metabolic and innate immune remodeling in the ileal mucosa.

**Figure 5.**
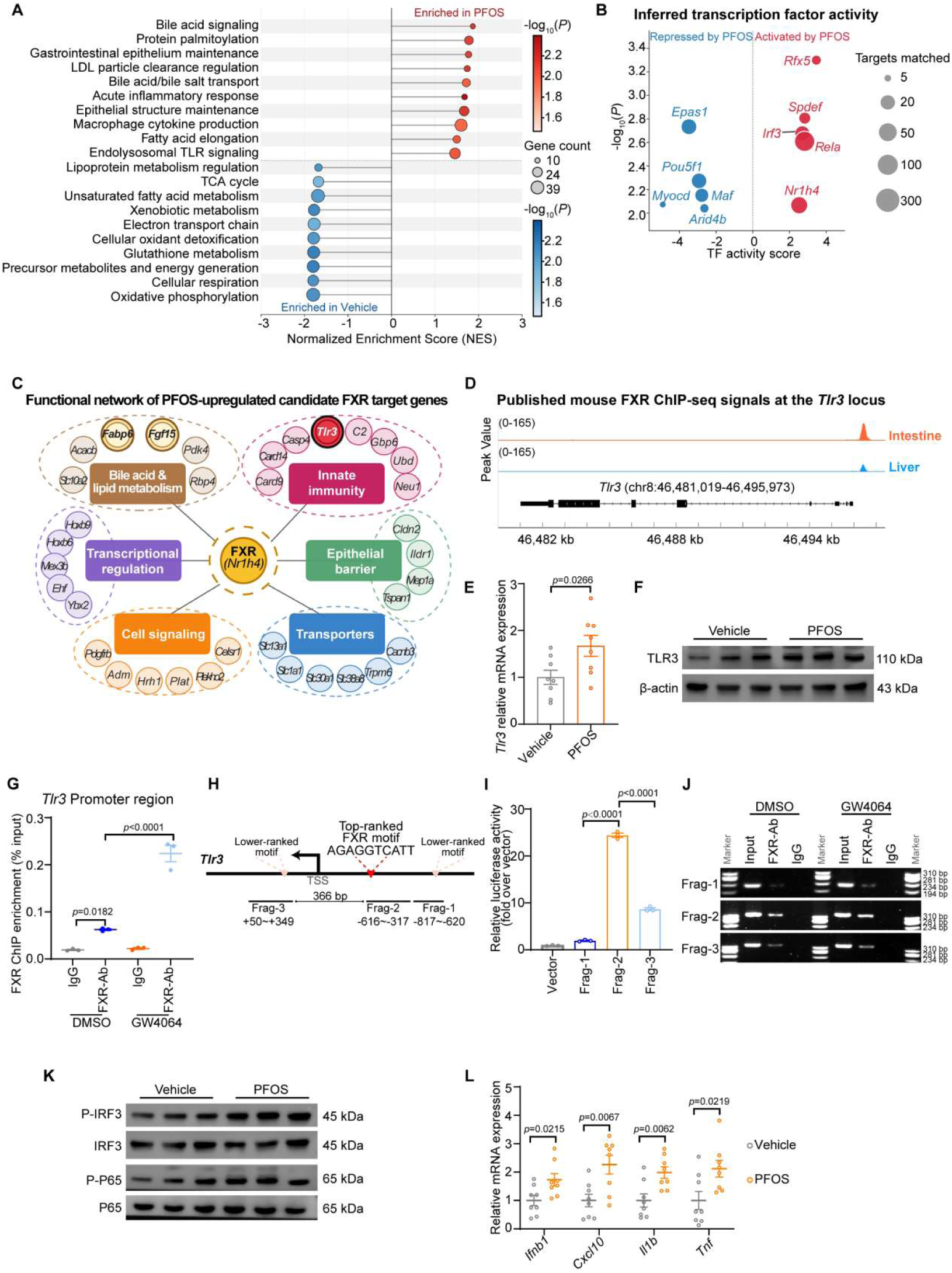
Tlr3 is a direct transcriptional target of intestinal FXR. **A**, Gene set enrichment analysis of ileal RNA-seq from high-cholesterol diet-fed *ApoE*⁻/⁻ mice exposed to Vehicle or PFOS. **B**, Inferred transcription factor activity in ileal tissues. **C**, Functional annotation of PFOS-upregulated candidate FXR target genes identified by integrating ileal RNA-seq with published intestinal FXR ChIP-seq data. **D**, Published FXR ChIP-seq signals at the *Tlr3* locus in mouse intestine and liver. **E** and **F**, *Tlr3* mRNA (**E**, n = 8 per group) and TLR3 protein (**F**, n = 3 per group) expression in ileal tissues. **G**, ChIP-qPCR analysis of FXR enrichment at the Tlr3 promoter in intestinal organoids treated with DMSO or GW4064. n = 3 per group. **H**, Schematic of predicted FXR-binding motifs and reporter fragments in the Tlr3 promoter. **I**, Dual-luciferase reporter assay of *Tlr3* promoter fragments. n = 3 per group. **J**, Agarose gel electrophoresis of PCR products from FXR ChIP-enriched DNA at the indicated *Tlr3* promoter fragments. **K**, Immunoblot analysis of IRF3 and p65 phosphorylation in ileal tissues. n=3 per group. **L**, qPCR analysis of ileal inflammatory and interferon-related genes. n = 8 per group. Data are presented as mean ± SEM. Each dot represents one biological replicate. *P* values are indicated in the panels. Two-tailed Student’s t test (**E**,**L**). One-way ANOVA with Tukey’s post hoc test (**G**,**I**). Source data are provided as a Source Data file.

**Supplementary Figure 5.**
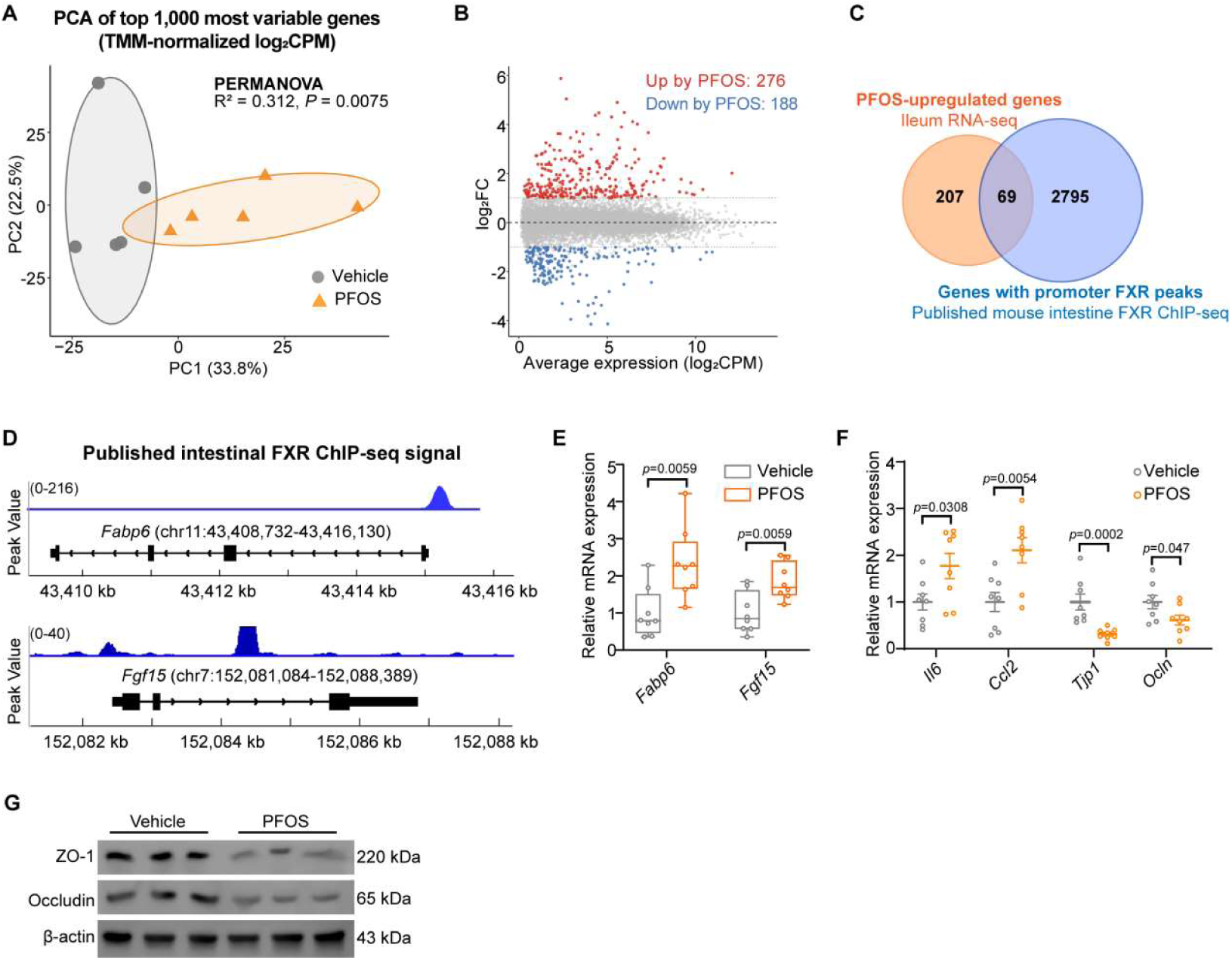
Tlr3 is a direct transcriptional target of intestinal FXR. Related to Figure5. **A**, PCA of the top 1,000 most variable genes based on TMM-normalized log₂CPM values from ileal RNA-seq of high-cholesterol diet-fed *ApoE*⁻/⁻ mice exposed to Vehicle or PFOS; group separation was assessed by PERMANOVA. **B**, Differential expression analysis of ileal RNA-seq; differentially expressed genes were defined as |log₂FC| > 1 and *P* < 0.05. **C**, Overlap between PFOS-upregulated genes and genes with promoter FXR peaks in published intestinal FXR ChIP-seq data. **D**, Published intestinal FXR ChIP-seq signals at the canonical FXR target loci *Fabp6* and *Fgf15*. **E**, qPCR validation of *Fabp6* and *Fgf15* expression in ileal tissues. n = 8 per group. **F**, qPCR analysis of ileal inflammatory and epithelial barrier-related genes. n = 8 per group. **G**, Immunoblot analysis of ZO-1 and Occludin in ileal tissues. n = 3 per group. Data are presented as mean ± SEM. Each dot represents one biological replicate. *P* values are indicated in the panels. Two-tailed Student’s t test (**E-F**). Two-sided Mann Whitney U test (**F**) Source data are provided as a Source Data file.

Transcription factor activity inferred using decoupleR and the CollecTRI network showed increased activities of FXR, IRF3, and RELA following PFOS exposure (Figure 5B). Because TUDCA and GUDCA have been identified as endogenous FXR antagonists, the reduction in TUDCA observed after PFOS exposure may have diminished antagonistic restraint on intestinal FXR signaling^[28]^. The parallel increases in inferred IRF3 and RELA activities further suggested coordinated activation of bile acid sensing and inflammatory signaling.

To identify potential direct FXR targets, we integrated published mouse intestinal FXR ChIP-seq data with PFOS-upregulated ileal genes. Genes were considered candidate targets if they contained an FXR-binding peak within −2 kb to +500 bp of the transcription start site and were upregulated after PFOS exposure. This analysis identified 69 overlapping genes (Supplementary Figure 5C). Functional annotation indicated that the major represented categories included bile acid and lipid metabolism, innate immunity, transcriptional regulation, epithelial barrier function, signaling, and transport (Figure 5C). The canonical FXR targets *Fabp6* and *Fgf15* showed promoter FXR-binding signals and increased ileal expression after PFOS exposure, supporting the validity of this integrative approach (Supplementary Figure 5D and E). Among the candidate targets, *Tlr3* was of particular interest because it encodes an endosomal pattern-recognition receptor that senses double-stranded RNA and activates downstream IRF3- and NF-κB-dependent inflammatory signaling.

Published FXR ChIP-seq data revealed a prominent FXR-binding signal at the *Tlr3* promoter, with stronger enrichment in the intestine than in the liver (Figure 5D). PFOS exposure also increased *Tlr3* mRNA and TLR3 protein expression in ileal tissues (Figure 5E and F). ChIP-qPCR in intestinal organoids showed basal FXR occupancy at the *Tlr3* promoter, which was further enhanced by the FXR agonist GW4064 (Figure 5G). JASPAR analysis identified three candidate FXR-binding regions near the *Tlr3* transcription start site (Figure 5H). Among the corresponding reporter constructs, Frag-2 (−616 to −317 bp) produced greater luciferase activity than Frag-1 or Frag-3 (both *P* < 0.0001), identifying this segment as the predominant FXR-responsive region (Figure 5I). Fragment-specific PCR using FXR ChIP-enriched DNA likewise showed the strongest amplification for Frag-2, with no obvious signal in the IgG control (Figure 5J). Together, these data identify the −616 to −317 bp region as the major FXR-bound and FXR-responsive region within the *Tlr3* promoter.

PFOS exposure was also accompanied by increased phosphorylation of p65 and IRF3 in ileal tissues (Figure 5K), together with upregulation of the inflammatory genes *Il1b*, *Tnf*, *Il6*, and *Ccl2* and the IRF3-associated genes *Ifnb1* and *Cxcl10* (Figure 5L and Supplementary Figure 5F). In parallel, *Tjp1* and *Ocln* expression and the corresponding ZO-1 and Occludin protein levels were reduced, indicating impaired epithelial barrier integrity (Supplementary Figure 5F and G). Collectively, these findings identify *Tlr3* as a direct transcriptional target of intestinal FXR and establish an intestinal FXR-TLR3 transcriptional link associated with activation of NF-κB/IRF3 signaling after PFOS exposure.

### Intestinal TLR3 partially mediates the proatherogenic effects of intestinal FXR

To determine whether intestinal FXR mediates the proatherogenic effects of PFOS, we generated mice with intestinal epithelial cell-specific deletion of *Fxr* (*Fxr*^ΔIE^) and used their *Fxr*^fl/fl^ littermates as controls. To induce atherosclerosis, both genotypes were administered AAV8-PCSK9^D377Y^, fed a HCD for 8 weeks, and exposed to PFOS for 5 weeks (n = 6 per group; Supplementary Figure 6A).

PFOS exposure activated intestinal FXR signaling in *Fxr*^fl/fl^ mice, whereas this activation was abolished in *Fxr*^ΔIE^ mice (Supplementary Figure 6B). Notably, Oil Red O staining showed that *Fxr*^ΔIE^ mice displayed markedly reduced atherosclerotic lesion areas in both the whole aorta and aortic root compared with *Fxr*^fl/fl^ mice, irrespective of PFOS exposure (Figure 6A-D). Consistently, macrophage infiltration within aortic root lesions was also lower in *Fxr*^ΔIE^ mice than in *Fxr*^fl/fl^ controls (Figure 6C,E). In parallel, circulating levels of IL-1β, TNF-α, and MCP-1 were significantly reduced in *Fxr*^ΔIE^ mice (Figure 6F-H). In addition, intestinal *Tlr3* mRNA expression was reduced in *Fxr*^ΔIE^ mice (Supplementary Figure 6C). Western blot analysis showed that PFOS-induced activation of intestinal TLR3 downstream signaling was markedly attenuated by intestinal FXR deletion, as evidenced by reduced phosphorylation of p65 and IRF3, with little change in total p65 or IRF3 protein abundance (Figure 6I). Notably, the suppression of p65 phosphorylation was accompanied by decreased intestinal expression of NF-κB-regulated pro-inflammatory genes, including *Il6*, *Il1b*, *Tnf*, and *Ccl2*, indicating that intestinal FXR deficiency blunted the PFOS-induced TLR3-NF-κB inflammatory response(Supplementary Figure 6E). Although IRF3-related genes, including *Ifnb1* and the interferon-stimulated gene *Cxcl10*, were also downregulated, the predominant effect was reflected by attenuation of the intestinal NF-κB inflammatory program(Supplementary Figure 6D). Concurrently, intestinal tight junction protein expression was restored in *Fxr*^ΔIE^ mice, suggesting preservation of epithelial barrier integrity ( Supplementary Figure 6F-G).Furthermore, hepatic and plasma levels of cholesterol and triglycerides were significantly reduced in *Fxr*^ΔIE^ mice (Figure 6J-K and Supplementary Figure 6H-I). Collectively, these findings suggest that intestinal FXR ablation mitigates PFOS-induced intestinal inflammation, barrier dysfunction, and hypercholesterolemia, thereby contributing to the improvement of atherosclerotic burden in this model.

**Figure 6.**
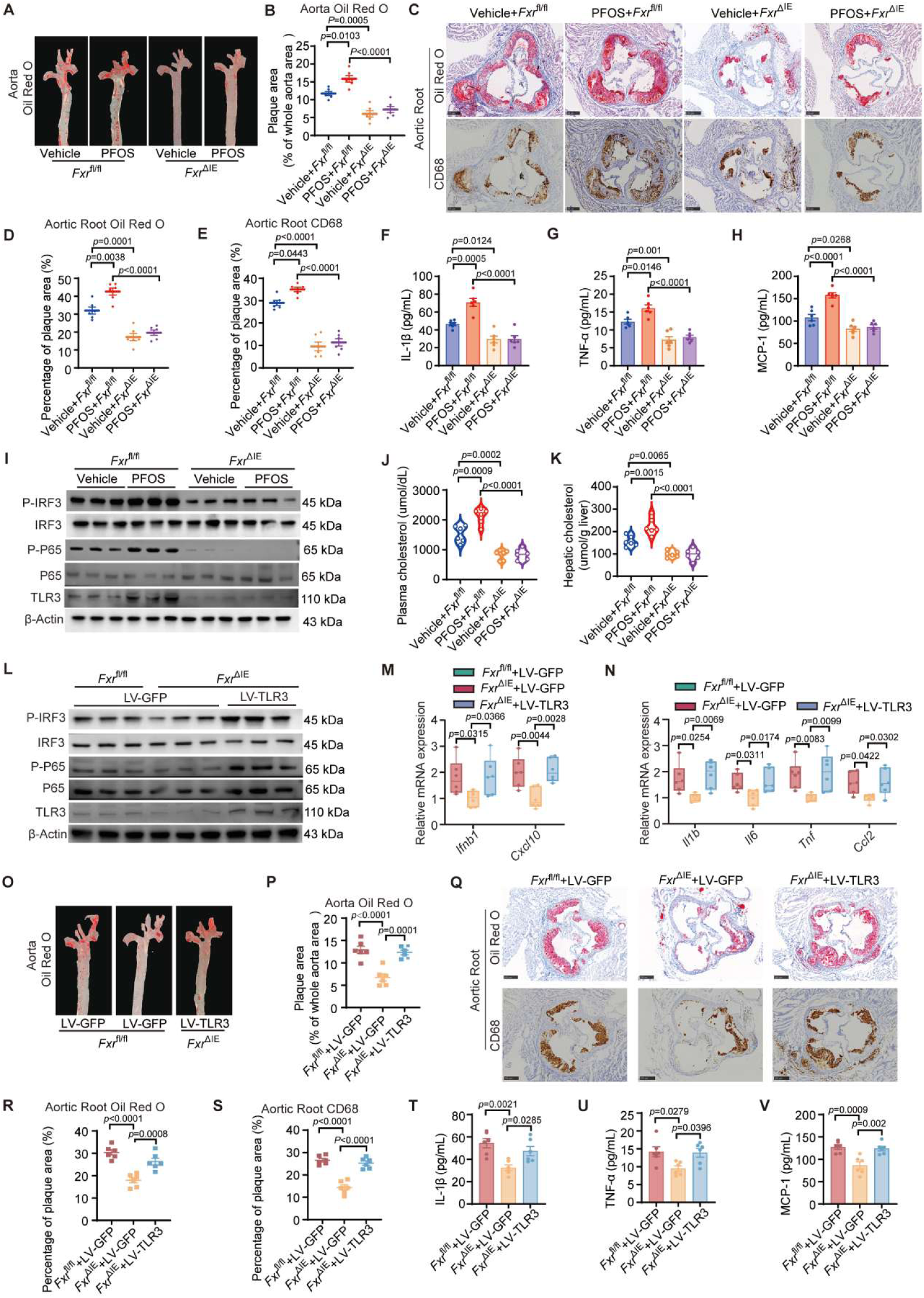
Intestinal TLR3 partially mediates the proatherogenic effects of intestinal FXR. **A** and **B**, Representative images of aortas stained with Oil Red O (**A**) and quantification of the lesions (**B**). n = 6 per group. **C**, Representative images of Oil Red O staining and CD68^+^ macrophage staining of aortic root cross-sections. Scale bar, 250 μm. n = 6 per group. **D** and **E**, Quantification analysis of Oil Red O staining (**D**) and CD68^+^ macrophage staining (**E**) of aortic roots. n = 6 per group. **F-H**, Levels of the proinflammatory cytokines IL-1β (**F**), TNF-α (**G**), and MCP-1 (**H**) in the plasma. n = 6 per group. **I**, Protein expression level of IRF3, p65 phosphorylation and TLR3. n = 3 per group. **J** and **K**, Plasma (**J**) and hepatic (**K**) cholesterol levels of the indicated mice.n = 8 per group. **L**, Protein expression level of IRF3, p65 phosphorylation and TLR3. n = 3 per group. **M** and **N**, mRNA levels of ileal inflammatory and interferon-related genes. n = 6 per group. **O** and **P**, Representative images of aortas stained with Oil Red O (**O**) and quantification of the lesions (**P**). n = 6 per group. **Q**, Representative images of Oil Red O staining and CD68^+^ macrophage staining of aortic root cross-sections. Scale bar, 250 μm. n = 6 per group. **R** and **S**, Quantification analysis of Oil Red O staining (**R**) and CD68^+^ macrophage staining (**S**) of aortic roots. n = 6 per group. **T-V**, Levels of the proinflammatory cytokines IL-1β (**T**), TNF-α (**U**), and MCP-1 (**V**) in the plasma. n = 6 per group. Data are presented as mean ± SEM. Each dot represents one biological replicate. *P* values are indicated in the panels. One-way ANOVA with Tukey’s post hoc test (**B**,**D-H**,**J-K**,**M-N**,**P**,**R-V**). Kruskal-Wallis with Dunn’s test (**N**) Source data are provided as a Source Data file.

**Supplementary Figure 6.**
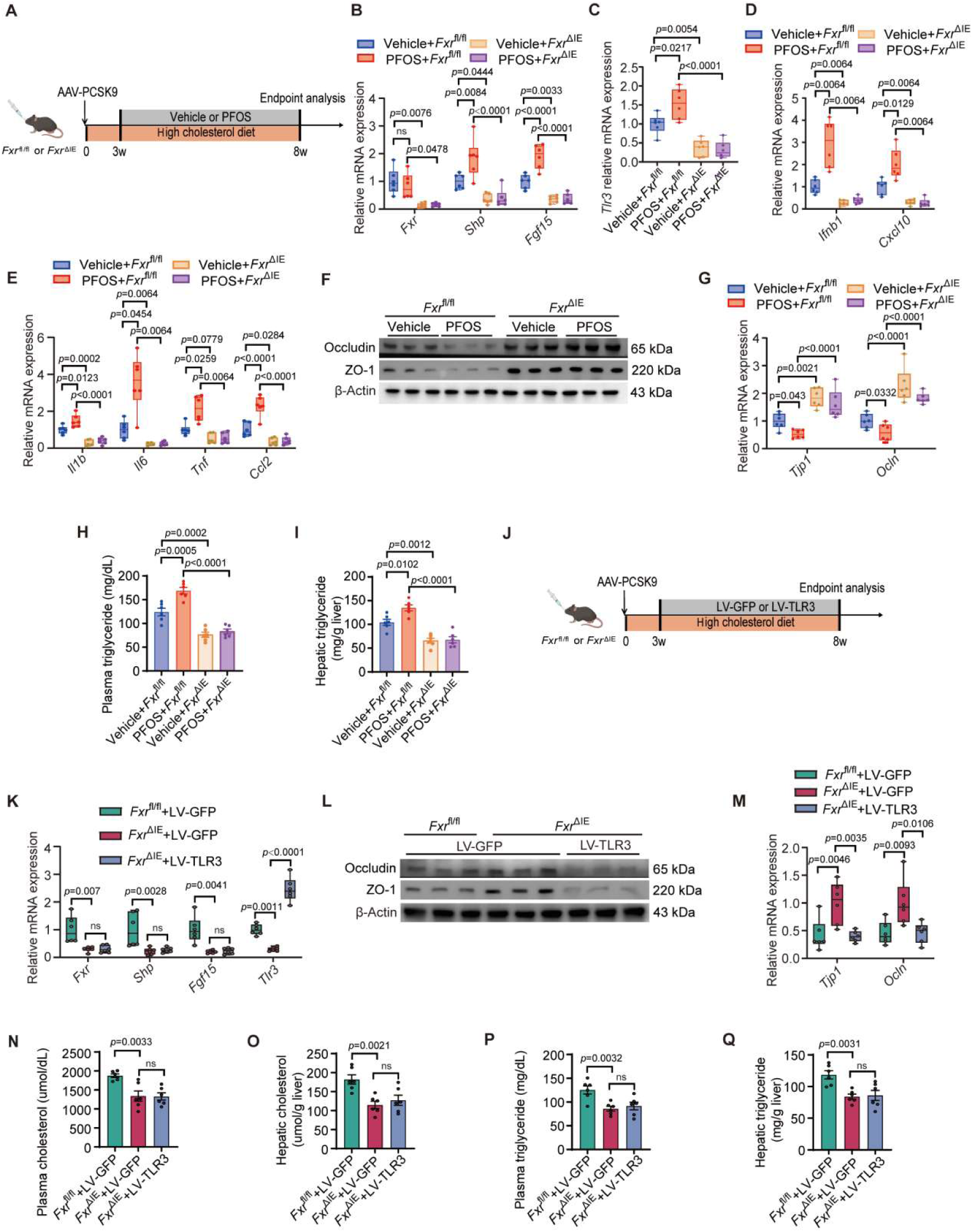
Intestinal TLR3 partially mediates the proatherogenic effects of intestinal FXR. Related to Figure 6. **A**, Schematic of the experimental design evaluating the role of FXR knockout in PFOS-aggravated atherosclerotic plaque formation. *Fxr*^fl/fl^ and *Fxr*^ΔIE^ mice were treated with AAV-PCSK9, a high-cholesterol diet, and PFOS or vehicle. n = 6 per group. **B**, *Fxr*, *Shp* and *Fgf15* mRNA levels in the ileum. n = 6 per group. **C**, mRNA levels of *Tlr3* in the ileum. n = 6 per group. **D** and **E**, mRNA levels of ileal inflammatory and interferon-related genes. n = 6 per group. **F** and **G**, ZO-1 and Occludin protein level (**F**, n = 3 per group) and mRNA levels (**G**, n = 6 per group). **H** and **I**, Plasma (**H**) and hepatic (**I**) triglyceride levels of the indicated mice. n = 6 per group. **J**, Schematic of the experimental design showing that intestinal TLR3 overexpression counteracts the atheroprotective effect of FXR deficiency. *Fxr*^fl/fl^ and *Fxr*^ΔIE^ mice were treated with AAV-PCSK9, a high-cholesterol diet, and LV-GFP or LV-TLR3. n = 6 per group. **K**, *Fxr*, *Shp*, *Fgf15* and *Tlr3* mRNA levels in the ileum.n = 6 per group. **L** and **M**, ZO-1 and Occludin protein level (**L**, n = 3 per group) and mRNA levels (**M**, n = 6 per group). **N-Q**, Plasma (**N**) and hepatic (**O**) cholesterol levels and plasma (**P**) and hepatic (**Q**) triglyceride levels of the indicated mice. n = 6 per group. Data are presented as mean ± SEM. Each dot represents one biological replicate. P values are indicated in the panels. Kruskal-Wallis with Dunn’s test (**B**, **K**). One-way ANOVA with Tukey’s post hoc test (**B-C**, **E**, **G-I**, **K**, **M-Q**). Mann-Whitney U test with the Bonferroni adjustment (**D-E**). Source data are provided as a Source Data file.

To determine whether TLR3 functions as a downstream effector of intestinal FXR, we restored intestinal TLR3 expression in *Fxr*^ΔIE^ mice by local delivery of LV-TLR3 and established atherosclerosis using AAV8-PCSK9^D377Y^ (Supplementary Figure 6J). LV-TLR3 selectively increased TLR3 expression in the ileum and reactivated downstream NF-κB and IRF3 signaling, as indicated by increased phosphorylation of p65 and IRF3 and elevated expression of associated inflammatory genes (Figure 6L-N and Supplementary Figure 6K). Moreover, intestinal TLR3 overexpression partially reversed the reduction in proinflammatory gene expression and the restoration of tight-junction proteins observed in *Fxr*^ΔIE^ mice, indicating that TLR3 reactivation compromises the improvement in intestinal inflammation and epithelial barrier integrity conferred by intestinal FXR deficiency (Supplementary Figure 6L-M).

With respect to the atherosclerotic phenotype, Oil Red O staining showed that TLR3 overexpression partially reversed the reduction in lesion area in both the whole aorta and aortic root of *Fxr*^ΔIE^ mice (Figure 6O-Q). Consistently, CD68 immunostaining revealed greater macrophage infiltration within aortic root plaques in LV-TLR3-treated *Fxr*^ΔIE^ mice than in LV-GFP-treated controls (Figure 6Q,S). In addition, forced TLR3 overexpression blunted the decrease in circulating IL-1β, TNF-α, and MCP-1 levels in *Fxr*^ΔIE^ mice (Figure 6T-V). Notably, LV-TLR3 treatment did not significantly affect cholesterol or triglyceride levels in the liver or plasma (Supplementary Figure 6N-Q). Collectively, these findings indicate that intestinal TLR3 functions as an important downstream effector of FXR in regulating intestinal inflammation and barrier integrity. Reactivation of TLR3 primarily enhances intestinal NF-κB/IRF3 inflammatory signaling, disrupts epithelial barrier integrity, and promotes systemic inflammation, thereby partially offsetting the atheroprotective effects of intestinal FXR deficiency independently of changes in circulating lipid levels.

### TUDCA and CU-CPT4a limit the progression of established atherosclerosis

To explore whether targeting the intestinal FXR-TLR3 axis could serve as a therapeutic strategy for established atherosclerotic lesions, we first evaluated the effect of exogenous TUDCA based on our previous findings that PFOS exposure reduced TUDCA levels and activated intestinal FXR-TLR3 signaling. After 5 weeks of PFOS exposure, mice were treated with TUDCA by oral gavage for an additional 3 weeks, followed by endpoint analysis (Supplementary Figure 7A). Oil Red O staining of the whole aorta and aortic root sections showed that TUDCA treatment significantly limited the further progression of established atherosclerotic lesions (Figure 7A-D). Consistently, CD68 immunostaining and plasma inflammatory cytokine measurements showed that TUDCA reduced macrophage accumulation within plaques and attenuated systemic inflammatory responses compared with PFOS-treated mice (Figure 7C,E,F and Supplementary Figure 7B and C).

**Figure 7.**
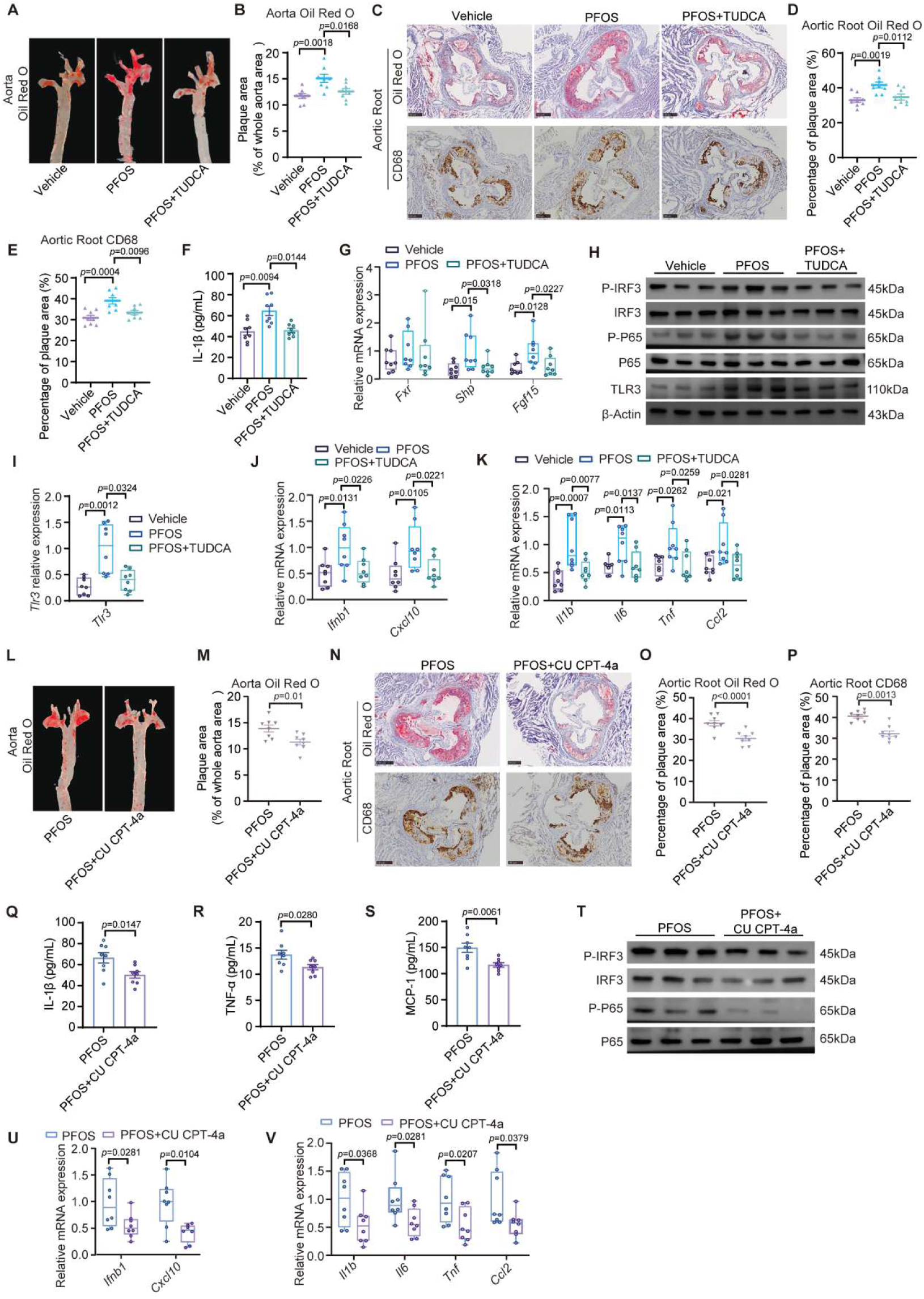
TUDCA and CU-CPT4a limit the progression of established atherosclerosis. **A** and **B**, Representative images of aortas stained with Oil Red O (**A**) and quantification of the lesions (**B**). n = 8 per group. **C**, Representative images of Oil Red O staining and CD68^+^ macrophage staining of aortic root cross-sections. Scale bar, 250 μm. n = 8 per group. **D** and **E**, Quantification analysis of Oil Red O staining (**D**) and CD68^+^ macrophage staining (**E**) of aortic roots. n = 8 per group. **F**, Levels of the proinflammatory cytokines IL-1βin the plasma. n = 8 per group. **G**, *Fxr*, *Shp* and *Fgf15* mRNA levels in the ileum. n = 8 per group. **H**, Protein expression level of IRF3, p65 phosphorylation and TLR3. n = 3 per group. **I**, mRNA levels of *Tlr3* in the ileum. n = 8 per group. **J** and **K**, mRNA levels of ileal inflammatory and interferon-related genes. n = 8 per group. **L** and **M**, Representative images of aortas stained with Oil Red O (**L**) and quantification of the lesions (**M**). n = 8 per group. **N**, Representative images of Oil Red O staining and CD68^+^ macrophage staining of aortic root cross-sections. Scale bar, 250 μm. n = 8 per group. **O** and **P**, Quantification analysis of Oil Red O staining (**O**) and CD68^+^ macrophage staining (**P**) of aortic roots. n = 8 per group. **Q-S**, Levels of the proinflammatory cytokines IL-1β (**Q**), TNF-α (**R**), and MCP-1 (**S**) in the plasma. n = 8 per group. **T**, Protein expression level of IRF3, p65 phosphorylation. n = 3 per group. **U** and **V**, mRNA levels of ileal inflammatory and interferon-related genes. n = 8 per group. Data are presented as mean ± SEM. Each dot represents one biological replicate. *P* values are indicated in the panels. One-way ANOVA with Tukey’s post hoc test (**B**,**D-G**,**J-K**). Kruskal-Wallis with Dunn’s test (**I**). Two-tailed Student’s t test (**M**,**O-S**). Two-sided Mann-Whitney U test (**U-V**). Source data are provided as a Source Data file.

**Supplementary Figure 7.**
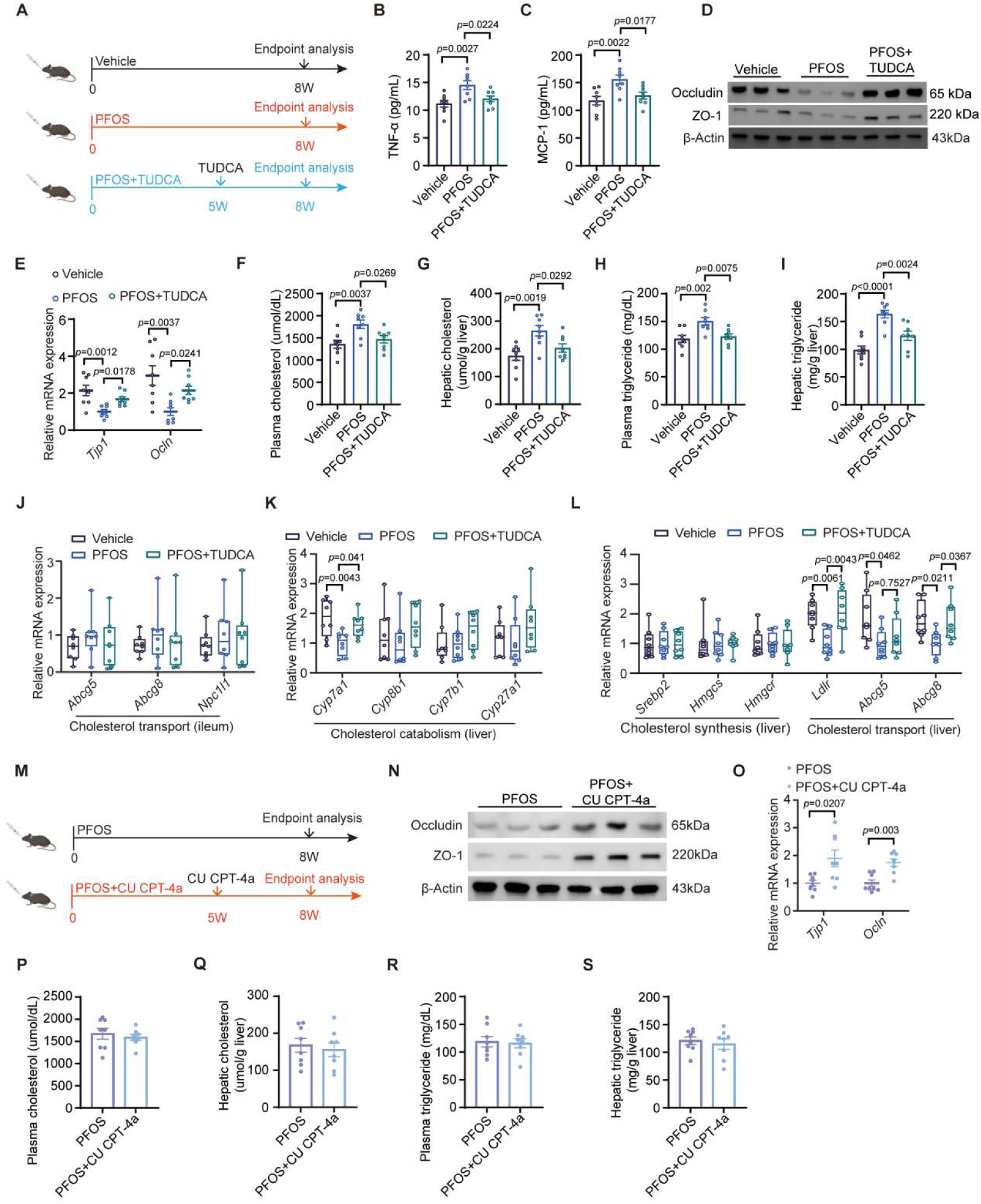
TUDCA and CU-CPT4a limit the progression of established atherosclerosis. Related to Figure 7. **A**, Schematic illustration of the TUDCA treatment protocol. n = 8 per group. **B** and **C**, Levels of the proinflammatory cytokines TNF-α (**B**) and MCP-1 (**C**) in the plasma. n = 8 per group. **D** and **E**, ZO-1 and Occludin protein level (**D**, n = 3 per group) and mRNA levels (**E**, n = 8 per group). **F**-**I**, Plasma (**F**) and hepatic (**G**) cholesterol levels and plasma (**H**) and hepatic (**I**) triglyceride levels of the indicated mice. n = 8 per group. **J**, Levels of mRNAs encoded by genes related to cholesterol transport in the ileum. n = 8 per group. **K** and **L**, Levels of mRNAs encoded by genes related to cholesterol elimination (**K**), cholesterol transport and cholesterol synthesis (**L**) in the liver. n = 8 per group. **M**, Schematic illustration of the CU CPT-4a treatment protocol. n = 8 per group. **N** and **O**, ZO-1 and Occludin protein level (**N**, n = 3 per group) and mRNA levels (**O**, n = 8 per group). **P-S**, Plasma (**p**) and hepatic (**Q**) cholesterol levels and plasma (**R**) and hepatic (**S**) triglyceride levels of the indicated mice. n = 8 per group. Data are presented as mean ± SEM. Each dot represents one biological replicate. *P* values are indicated in the panels. One-way ANOVA with Tukey’s post hoc test (**B-C**, **F-I**, **K-L**). Kruskal-Wallis with Dunn’s test (**E**, **J**). Two-sided Mann-Whitney U test (**O**). Two-tailed Student’s t test (**P-S**). Source data are provided as a Source Data file.

We next examined ileal FXR signaling and its downstream TLR3 inflammatory pathway. Compared with PFOS-treated mice without TUDCA intervention, TUDCA treatment suppressed intestinal FXR signaling activity (Figure 7G). Moreover, TUDCA reduced the expression of the FXR target gene *Tlr3* and inhibited activation of TLR3 downstream NF-κB and IRF3 signaling, as reflected by decreased phosphorylation of p65 and IRF3 and reduced expression of NF-κB/IRF3-associated inflammatory genes (Figure 7H-K). In parallel with the attenuation of inflammatory signaling, TUDCA restored intestinal tight junction protein expression, suggesting improved epithelial barrier integrity (Supplementary Figure 7D and E). In addition, TUDCA treatment reduced plasma and hepatic cholesterol and triglyceride levels (Supplementary Figure 7F-I). Although the mRNA expression of ileal cholesterol transport genes remained largely unchanged (Supplementary Figure 7J), hepatic *Cyp7a1*, *Ldlr*, and *Abcg8* expression was increased after TUDCA treatment (Supplementary Figure 7K-L). These findings suggest that TUDCA limits atherosclerotic lesion progression by suppressing the intestinal FXR-TLR3 inflammatory axis and improving cholesterol metabolism.

To further assess the therapeutic potential of TLR3 as a downstream target, PFOS-exposed mice were treated daily with the TLR3 inhibitor CU CPT-4a by oral gavage for 3 weeks after 5 weeks of PFOS exposure (Supplementary Figure 7M). CU CPT-4a treatment reduced atherosclerotic lesion areas in both the whole aorta and aortic root (Figure 7L-O). Macrophage accumulation within aortic root plaques was also lower in CU CPT-4a-treated mice than in PFOS-treated controls (Figure 7N,P). In addition, plasma IL-1β, TNF-α, and MCP-1 levels were significantly decreased after CU CPT-4a treatment (Figure 7Q-S).

As expected, oral CU CPT-4a markedly inhibited PFOS-induced activation of intestinal TLR3 downstream signaling, as evidenced by reduced phosphorylation of p65 and IRF3, together with decreased expression of NF-κB/IRF3-associated inflammatory genes (Figure 7T-V). Meanwhile, intestinal tight junction protein expression was also restored following CU-CPT4a treatment (Supplementary Figure 7N and O).These results indicate that CU CPT-4a primarily suppresses TLR3-mediated NF-κB/IRF3 inflammatory signaling, thereby reducing intestinal inflammatory gene expression and systemic inflammation. Notably, CU CPT-4a treatment did not significantly alter cholesterol or triglyceride levels in the plasma or liver (Supplementary Figure 7P-S), suggesting that its atheroprotective effect is mainly mediated by inhibition of inflammatory signaling rather than changes in lipid metabolism.

Collectively, these data demonstrate that TUDCA acts upstream by suppressing the intestinal FXR-TLR3 axis and improving inflammation, barrier integrity, and lipid metabolism, whereas CU CPT-4a acts downstream by directly inhibiting TLR3-NF-κB/IRF3 inflammatory signaling. Both interventions attenuate the progression of established atherosclerotic lesions, supporting the intestinal FXR-TLR3 inflammatory axis as a therapeutically targetable pathway in atherosclerosis.

## Discussion

In this study, we identify the gut microbiota as a causal mediator of PFOS-exacerbated atherosclerosis and reveal how bacterial adaptation to chemical stress is translated into host immunometabolic dysfunction. The gut microbiota-bile acid–intestinal FXR–TLR3 axis identified here extends the understanding of PFOS cardiovascular toxicity beyond direct host injury and positions the intestinal microbiota as an active mediator linking environmental exposure to systemic vascular inflammation.

Microbiota depletion markedly attenuated the vascular, inflammatory, and metabolic effects of PFOS, whereas microbiota from PFOS-exposed donors transferred these abnormalities to recipient mice. These findings do not exclude direct host effects of PFOS, but demonstrate that microbial remodeling is required for the full proatherogenic response. Among the altered taxa, viable *B.caecimuris* aggravated HCD-induced atherosclerosis in the absence of direct PFOS exposure, indicating that active microbial functions contribute to its pathogenic effect. This observation is consistent with the broader concept that pollutant-induced changes in microbial composition and function can mediate systemic metabolic and inflammatory consequences ^[5,15,19–22]^.

Our data further suggest that TolC-mediated efflux enables *B.caecimuris* to better tolerate PFOS and maintain a higher intestinal abundance during exposure. TolC is the outer-membrane component of tripartite efflux systems in Gram-negative bacteria and participates in the export of structurally diverse antimicrobial agents, bile salts, and toxic compounds ^[29,30]^. Such efflux systems contribute not only to antimicrobial resistance but also to bacterial stress adaptation, ecological fitness, and persistence in hostile environments ^[29–31]^. TolC therefore appears to act primarily as a bacterial fitness determinant rather than as a direct regulator of bile acid metabolism. More broadly, these findings support the concept that persistent environmental chemicals can act as ecological selection pressures that favor microorganisms equipped with xenobiotic-tolerance mechanisms, thereby shifting the gut microbiota toward disease-promoting functional states ^[19,20]^.

Bile acid remodeling provides the metabolic link between microbial selection and host signaling. The gut microbiota extensively modifies the size and composition of the bile acid pool through deconjugation and subsequent biotransformation, thereby influencing host metabolic and immune signaling ^[16,32]^. PFOS exposure reduced TUDCA in ileal contents and plasma, while viable *B.caecimuris* reproduced this alteration in vivo and depleted TUDCA in vitro. Although other microbial and host pathways may also contribute, these results support an important role for *B.caecimuris* in PFOS-associated TUDCA loss. Ileal transcriptomic analysis further indicated enrichment of bile acid-related pathways and increased inferred FXR activity after PFOS exposure. Because TUDCA and GUDCA have been identified as endogenous FXR antagonists ^[28]^, reduced TUDCA may relieve antagonistic restraint within the local bile acid pool and thereby contribute to enhanced intestinal FXR signaling.

A major advance of this study is the identification of Tlr3 as a direct transcriptional target of intestinal FXR. TLR3 is an endosomal pattern-recognition receptor that recognizes double-stranded RNA and signals exclusively through the adaptor TRIF to activate NF-κB- and IRF3-dependent inflammatory and type I interferon responses ^[33,34]^. By increasing TLR3 abundance, intestinal FXR connects bile acid sensing to innate immune activation and may increase mucosal responsiveness to nucleic acid-associated stimuli. This finding broadens the conventional view of FXR from a regulator of bile acid and lipid metabolism to a transcriptional modulator of innate immune sensing.

The functional hierarchy of this pathway was supported by genetic and pharmacological interventions. Intestinal FXR deletion reduced inflammation, circulating lipid levels, and atherosclerotic burden, whereas restoration of intestinal TLR3 reactivated NF-κB/IRF3 signaling and partially reversed vascular protection without restoring the lipid phenotype. These findings identify TLR3 as an important, but not exclusive, downstream effector of intestinal FXR and distinguish a relatively lipid-independent inflammatory branch from the broader metabolic actions of FXR. Consistently, TUDCA improved both metabolic and inflammatory abnormalities, whereas CU-CPT4a primarily suppressed downstream inflammatory signaling.

These results also highlight the context-dependent biology of FXR. FXR regulates bile acid, lipid, and glucose homeostasis and can exert anti-inflammatory effects in hepatic and immune-cell settings ^[35,36]^. However, intestinal FXR can also promote disease through tissue-specific metabolic and epithelial programs. Inhibition of the intestinal FXR-SMPD3-ceramide pathway has been shown to reduce atherosclerosis, demonstrating that intestinal FXR can exert proatherogenic metabolic effects ^[37]^. More recently, intestinal FXR activation was reported to promote epithelial ferroptosis, barrier dysfunction, and ILC3 impairment in intestinal disease ^[38]^. Our study adds a distinct mechanism by showing that intestinal FXR directly regulates a pattern-recognition receptor, thereby linking metabolic sensing to innate immune activation. The biological consequences of FXR activation therefore depend on the tissue, cell type, downstream transcriptional program, and disease context.

Several limitations should be acknowledged. First, the human analysis was observational and based on a modest sample size, particularly for participants with coronary angiography data. The association between fecal PFOS levels and coronary atherosclerotic burden therefore requires validation in larger, independent prospective cohorts. Second, although TUDCA was the most consistent bile acid alteration and was functionally validated, additional microbiota-derived signals may contribute to PFOS-induced host responses. Moreover, the endogenous or microbial nucleic acid ligands that activate the increased intestinal TLR3 pool remain to be identified.

In summary, this study expands the understanding of PFOS cardiovascular toxicity from direct host injury to microbiota-dependent immunometabolic reprogramming. Pollutant-tolerance mechanisms can reshape the intestinal ecosystem, alter microbial metabolism, and amplify host innate immune signaling. Microbial fitness, bile acid signaling, and the intestinal FXR-TLR3 inflammatory pathway may therefore represent interconnected targets for mitigating pollutant-associated cardiovascular risk.

## Methods

### Study population and sample collection

This study was conducted in the Department of Cardiology, Second Hospital of Tianjin Medical University. Participants were consecutively recruited from December 19, 2025. A total of 127 participants with paired serum and fecal samples were included in the present analysis. Serum and fecal samples were collected for the measurement of PFOS levels. Clinical characteristics, lipid-related biochemical indicators, and cardiovascular risk-related covariates were also collected. Among these participants, 82 underwent coronary angiography and had available GS score data for the assessment of coronary atherosclerotic burden. The study protocol was approved by the Ethics Committee of the Second Hospital of Tianjin Medical University (ky2025100), and all participants provided written informed consent.

### Measurement of lipid-related indicators

Lipid-related biochemical indicators were measured using a Mindray BS-2800M automatic biochemical analyzer with matched original reagents. Total cholesterol (TC) was measured using the CHOD-PAP enzymatic method. Triglycerides (TG) were measured using the GPO-PAP enzymatic method. High-density lipoprotein cholesterol (HDL-C) and low-density lipoprotein cholesterol (LDL-C) were measured using direct homogeneous assays. Additional lipid-related indicators, including very-low-density lipoprotein cholesterol (VLDL-C), apolipoprotein A1 (ApoA1), apolipoprotein B (ApoB), ApoA1/B, lipoprotein(a) [Lp(a)], and atherogenic index (AI), were obtained from clinical laboratory records.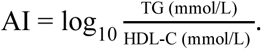.

### Coronary angiography and assessment of coronary atherosclerotic burden

Coronary angiography and quantitative coronary angiography (QCA) were performed according to standard clinical procedures. After entering the catheterization laboratory, patients were placed in the supine position on a 1000-mA angiography system. Following routine disinfection and draping, local anesthesia was administered using 1% lidocaine. After successful puncture of the right or left radial artery, a 6F sheath was inserted, and coronary angiography was performed using a 5F Tiger catheter or 6F Judkins and Amplatz catheters.

All coronary angiographic images were reviewed and recorded by trained members of the coronary interventional data analysis core team using QCA imaging analysis software (QAngio version 7.3; Medis Medical Imaging Systems). The analyzed parameters included the location of stenotic vessels, including the left main coronary artery, left anterior descending artery, left circumflex artery, and right coronary artery, as well as minimum lumen diameter, reference vessel diameter, percent diameter stenosis, and lesion length. Percent diameter stenosis was calculated as follows: diameter stenosis (%) = [1 - (minimum lumen diameter / reference vessel diameter)] × 100%.

Among participants with available coronary angiography data, the GS score was calculated to quantify the overall burden of coronary atherosclerosis. Analyses involving GS score were restricted to the 82 participants with complete coronary angiography-derived GS score data.

### PFOS Detection

Fecal samples: Freeze-dried fecal samples stored at -80 °C were thawed at 4 °C. Approximately 10 mg of sample was accurately weighed into a 1.5 mL EP tube, followed by the addition of 100 µL of water. The sample was homogenized at 3000 rpm for 100 s using a homogenizer with three 3 mm grinding beads. Then, 10 µL of 2.5 µg/mL internal standard solution and 300 µL of cold protein precipitation solution were added. After vortex mixing, the mixture was centrifuged at 16,000 g for 10 min, and 100 µL of the supernatant was collected. The extract was evaporated to dryness and reconstituted in methanol, followed by vortexing for 30 s. After centrifugation at 4 °C and 2000 g for 5 min, 20 µL of the supernatant was collected, diluted five-fold with methanol:water (8:2), and 5 µL was injected for analysis.

Serum samples: Serum samples stored at −80 °C were thawed at 4 °C. A 50 µL aliquot was transferred into a 1.5 mL EP tube, followed by the addition of 10 µL of 0.5 µg/mL internal standard solution and 300 µL of cold protein precipitation solution. After vortex mixing, the mixture was centrifuged at 16,000 g for 10 min, and 200 µL of the supernatant was collected. The extract was evaporated to dryness and reconstituted in methanol, followed by vortexing for 30 s. After centrifugation at 4 °C and 16,000 g for 5 min, the supernatant was collected, and 5 µL was injected for analysis.

Serum PFOS and fecal PFOS concentrations were measured from paired serum and fecal samples. Because PFOS concentrations showed skewed distributions, PFOS variables were log-transformed before statistical analysis. Serum and fecal PFOS were analyzed separately to evaluate their associations with lipid-related indicators and coronary atherosclerotic burden.

### Mouse models

SPF *ApoE*^-/-^ mice were obtained from VITAL RIVER (Beijing, China), and SPF intestinal-specific Fxr knockout (*Fxr^ΔIE^*) mice were kindly provided by Dr. Changtao Jiang. All mice were housed in a specific pathogen free (SPF) environment. 8 weeks old *ApoE*^-/-^ mice were housed at 23 ± 1 ℃, humidity (45 - 55%) on a 12 h light/12 h dark cycle and had free access to food and water. In animal experiments, 8 weeks old male mice were randomly assigned to different groups, and no mice were excluded from the analysis. All procedures were approved by the Institutional Animal Care and Use Committee (IACUC, protocol ZXHK-DWLL-2026-0080).

Plasma was collected to determine bile acid levels and inflammatory cytokine levels. The heart and vasculature were fixed in 4% paraformaldehyde. The liver, feces, cecum, and ileum were rapidly frozen and stored at −80 °C.

### Assessment of atherosclerotic lesions

Imaging of arterial lesions in vivo was performed with a high-frequency ultrasound system (Vevo 2100, Visualsonics) equipped with a linear array transducer (MS 550D, 22-55 MHz). Briefly, *ApoE*^−/−^ mice anesthetized with isoflurane were placed on a heated procedural board to maintain 37 °C and limbs were taped to electrocardiogram electrodes coated with electrode cream. The fur at the imaging location was shaved and warm ultrasound gel was liberally applied to ensure optimal image quality. The aortic sinus was imaged and visualized using a long-axis view, a CINE loop of 100 frames was stored for later off-line analysis and the time gain compensation curve was adjusted to produce uniform echo intensity. The gain was set to 30 dB and the dynamic range to 65 dB. Measurements were repeated three times at each site. After perfusion with PBS, the entire aorta and heart were harvested. The tissues were fixed in 10% neutral buffered formalin for 6 h, followed by dehydration in 20% sucrose solution overnight. Cardiac tissues were embedded in OCT compound for preparation of cryosections. Subsequently, the entire aorta was stained with Oil Red O and quantitatively analyzed using ImageJ software. Frozen sections were subjected to histological analysis following Oil Red O and CD68 staining. Quantitative analysis was performed using ImageJ software.

### Biochemical and immunological assays

To assess cholesterol and triglyceride levels, cholesterol assay kits (Sangon Biotech) and triglyceride assay kits (Sangon Biotech) were used to measure concentrations in plasma or liver samples. According to the manufacturer’s instructions, liver tissues were homogenized in cholesterol and triglyceride extraction buffer, and the supernatants were collected for measurement of cholesterol and triglyceride levels, which were then normalized to liver tissue weight. Plasma inflammatory cytokine levels were determined using Mouse IL-1β ELISA Kit, Mouse MCP-1 ELISA Kit, and Mouse TNF-α ELISA Kit (Sangon Biotech).

### Bacterial strains and culture

To confirm the species identity, *B.caecimuris* DSM26085 (purchased from BeNa Culture Collection, BNCC) was anaerobically cultured in brain heart infusion broth at 37 °C for 48-72 hours. Genomic DNA was subsequently extracted, and the 16S rRNA gene was amplified via PCR using primers 27F and 1492R. The amplified 16S gene was sequenced, and the resulting sequence was compared with the NCBI database to confirm species identity.

Cell pellets were collected by centrifugation at 8,000 × g for 10 min at 4°C, and oxygen-free PBS was used to resuspend cells to a final cell density of 1 × 10^9^ colony-forming units CFU)/mL. The cell suspension (200 μL) was then used for the animals’ oral administration every other day.The information on the chemical and biomolecular materials used in this study is provided in Supplementary Table S2.

### Minimum inhibitory concentration of *B.caecimuris* towards PFOS

To evaluate the susceptibility of *B.caecimuris* to PFOS, a minimum inhibitory concentration (MIC) test was performed. Briefly, two-fold serial dilutions of PFOS were prepared in a microtiter plate, followed by the addition of 100 µL of bacterial suspension to each well. The MIC was determined by measuring the absorbance at 600 nm (OD600) after incubation. Positive and negative controls were included for quality assurance.

### Ethidium Bromide-agar Cartwheel Method

Bacterial strains were inoculated into 5 mL of appropriate liquid culture medium and incubated until the optical density at 600 nm (OD600) reached 0.5. Agar plates containing ethidium bromide (EtBr) at concentrations ranging from 0 to 12 mg/L were prepared. Each plate was divided radially into up to 4 equal sectors. Using a sterile swab, the bacterial culture was inoculated onto the EtBr-supplemented agar plates, starting from the centre and spreading outward toward the periphery. The plates were incubated at 37°C for 48 hours and then examined under an appropriate ultraviolet (UV) light source, such as a handheld UV lamp. One reference strain was included on each plate as a comparative control. Relative EtBr efflux capacities of the bacterial strains were quantified against the reference strain using the following equation: index=MC_EtBr_(target strain)-MC_EtBr_(REF)/MC_EtBr_(REF), where MC_EtBr_(target strain) and MC_EtBr_(REF) correspond to the minimum EtBr concentrations necessary to produce a measurable fluorescent signal from the bacterial cell masses of the target strain and reference strains, respectively.

### Gene manipulation of *B. caecimuris*

The *tolC* family protein gene (designated *tolC*) was genetically modified using the pLGB31 vector. Briefly, approximately 1000 bp fragments corresponding to the upstream and downstream regions of *tolC*, each flanked by *XmaI* and *ClaI* restriction sites, were synthesised and assembled into the pUC57 vector. The pLGB31 plasmid was obtained from Addgene. For restriction digestion, 4 µg of each construct (pUC57 and pLGB31) were combined with 1 unit each of *XmaI* (NEB, UK) and *ClaI* (NEB, UK) in a total volume of 50 µL and incubated overnight at 37 °C. The resulting DNA fragments were separated via gel electrophoresis, and the bands of interest were excised using a sterile scalpel. These excised fragments were then purified using a commercial gel extraction kit. Subsequently, the purified inserts were ligated into the digested pLGB31 backbone using T4 DNA ligase (NEB, UK), following the manufacturer’s protocol. Putative positive clones were identified by PCR and confirmed by sequencing.

*Escherichia coli* strains were grown aerobically at 37 °C in LB medium, supplemented with 100 μg/mL carbenicillin when required. Plasmid pLGB31 was extracted from *E. coli* using a TIANGEN kit. *B.caecimuris* was cultured anaerobically at 37 °C on brain heart infusion (BHI) agar supplemented with 5 mg/L hemin and 2.5 μg/L vitamin K₁ (designated BHIS). Screening was performed on M9S plates, which contained M9 minimal medium supplemented with 50 mg/L L-cysteine, 5 mg/L hemin, 2.5 μg/L vitamin K₁, 2 mg/L FeSO₄·7H₂O, and 5 μg/L vitamin B₁₂. Inulin from chicory (Sigma-Aldrich) was added as a carbon source at 0.25% (w/v) to generate M9S-inulin plates.

Due to the inability of *B.caecimuris* to survive in the presence of *E. coli*, plasmid delivery was achieved via electroporation. Positive clone selection followed the method described by Leonor García-Bayona et al ^[39]^. In brief, a 7 mL culture of *B.caecimuris* was grown to an OD600 of 0.1-0.2. Cells were harvested by centrifugation (3,000×g, 10 min) and washed twice with 7 mL of 10% glycerol. The cell pellet was then resuspended in 100 µL of 10% glycerol and transferred to an electroporation cuvette. Two micrograms of pLGB31 plasmid DNA were added, and electroporation was performed at 12.5 kV/cm with a pulse duration of 5 ms ^[40]^. The electroporated cells were spread onto selective M9S-inulin plates. After 2-4 days of anaerobic incubation, single colonies were picked and screened by PCR to confirm cointegration. Following overnight growth, the cultures were diluted and plated (volumes of 50, 5, 1, and 0.2 µL) onto plates containing 10 mM rhamnose. Single colonies that appeared were picked and again verified by PCR.Genetic complementation of the *tolC* was performed using the intact coding region along with its upstream and downstream flanking sequences, as described above.

### DNA extraction and quantification of the abundance of *B.caecimuris* in feces

Fecal DNA was extracted from mouse fecal samples using the TIANamp Stool DNA Kit (TIANGEN) according to the manufacturer’s instructions. The concentration of fecal DNA was measured using a NanoDrop ND-2000 spectrophotometer (Thermo Scientific). Quantitative PCR (qPCR) was performed using a QuantStudio Dx Real-Time PCR system (Applied Biosystems) and ChamQ Universal SYBR qPCR Master Mix (Vazyme). A standard curve was generated using serially diluted genomic DNA from cultured *B.caecimuris.* The abundance of B. caecimuris was expressed as log10 copies per gram of fecal DNA. Primer sequences used for qPCR are listed in Table S3.

### Mouse experiment

In the PFOS intervention study, 8-week-old *ApoE*^-/-^ mice (male, n = 8 per group) were acclimatized for 1 week prior to experimentation. Mice were randomly assigned to two groups based on PFOS exposure status. All mice were fed a high-cholesterol diet (HCD; Research Diets, D12109C) to induce atherosclerosis. Based on previously reported dosing regimens ^[14,41–42]^, mice in the PFOS exposure group received daily oral gavage of PFOS (1 mg/kg body weight, dissolved in water) for 5 weeks. The gavage dose was adjusted according to body weight. Control mice received an equal volume of water by oral gavage once daily.

In the antibiotic treatment study, the antibiotic mixture was added to the drinking water of SPF 8-week-old *ApoE^-/-^* mice (male, n=6 per group) and continued for seven days.The broad-spectrum antibiotic combination consisted of vancomycin (50 mg/kg, Sigma), metronidazole (100 mg/kg, Sigma), kanamycin (100 mg/kg, Sigma), and ampicillin (100 mg/kg, Sigma) ^[43–44]^. Mice were then treated with the PFOS intervention and superimposed on a HCD for 5 weeks.

In the FMT study, recipient mice (8-week-old, male, n=8 per group) were pretreated with antibiotics as described above. Feces (20mg) were collected from the cecum of vehicle or PFOS-exposed mice, homogenized in pre-diluted PBS (1 mL per sample), shaken for 3 min, and then centrifuged at 4 ℃ for 3 min. Then 200 µl of supernatant was collected. A suspension of 100 microliters of the precipitate was given to recipient mice by oral gavage.

In the TUDCA and CU CPT-4a treatment experiment, 8-week-old *ApoE*^-/-^ mice (male, n = 8 per group) were first exposed to PFOS for 5 weeks and then randomly divided into groups and further treated with vehicle, TUDCA (50 mg/kg/day), or CU CPT-4a (5 mg/kg/day) for an additional 3 weeks, while being continuously fed a high-cholesterol diet daily.

In the *B.caecimuris* administration experiment, the strain was obtained from China BNBIO (BNCC). Its identity was confirmed by comparing 16S rRNA gene amplification sequences (27F and 1492R) with the NCBI reference database. *B.caecimuris* was cultured in Brain Heart Infusion (BHI) broth under anaerobic conditions at 37 °C for 24-48 h. Bacterial pellets were collected by centrifugation at 8,000 × g for 10 min at 4 °C and resuspended in anaerobic PBS to a final concentration of 1 × 10^8 CFU/mL. A 200 μL bacterial suspension was administered to mice by oral gavage every other day.Eight-week-old *ApoE*^-/-^ mice (male, n = 8 per group) were randomly divided into three groups and received oral gavage of water, heat-killed *B.caecimuris* (1 × 10^8 CFU/mouse), or live *B.caecimuris* (1 × 10^8 CFU/mouse), respectively. Mice were gavaged every other day for 5 weeks while continuously receiving a high-cholesterol diet.

Eight-week-old male *Fxr^fl/fl^* and *Fxr^ΔIE^* mice (n = 6 per group) were injected with AAV8-PCSK9(D377Y) via the tail vein, fed a high-cholesterol diet (HCD) for 8 weeks, and exposed to PFOS (1 mg/kg) for 5 weeks to assess the critical role of intestinal FXR in PFOS-exacerbated atherosclerosis.

Eight-week-old male *Fxr^fl/fl^* and *Fxr^ΔIE^* mice (n = 6 per group) were injected with AAV8-PCSK9(D377Y) via the tail vein and fed a high-cholesterol diet (HCD) for 8 weeks. After 3 weeks, mice received small-intestinal lentiviral injection to overexpress TLR3 in the ileum ^[45]^. Briefly, a 6-8 cm intestinal segment distal to the cecum was isolated, and both ends were ligated with clamps to prevent viral leakage and reflux of intestinal contents. A 3 mm longitudinal incision was made in the intestinal segment, and the segment was flushed with saline through an insulin syringe at a site 6 cm from the cecum. Subsequently, 0.2 mL of lentivirus expressing a scrambled control sequence or HBLV-m-TLR3 (Hanbio Biotechnology) was injected using an insulin syringe. After 20 min, the intestinal segment was flushed with saline, the clamps were removed, and the incision was closed with 10-0 sutures.

In the experiment using *B.caecimuris tolC* knockout and complementation strains, mice were orally administered the wild-type strain (10^8 CFU/mouse), the *tolC* mutant strain (10^8 CFU/mouse), or the complemented strain (10^8 CFU/mouse). All *ApoE*^−/−^ mice (8-week-old males, n = 6 per group) received PFOS (1 mg/kg body weight) by oral gavage for 2 consecutive weeks.

### Organoid culture

Small intestine tissues were isolated from mice and dissected, then washed 10 times with Dulbecco’s PBS (DPBS). The intestinal segments were subsequently incubated in Gentle Cell Dissociation Reagent (STEMCELL Technologies) to separate crypts and villi from the intestinal basement membrane. After centrifugation, isolated crypts were resuspended at a density of 6,000 crypts/mL in a 1:1 mixture of Matrigel (Corning) and IntestiCult organoid growth medium (OGM; STEMCELL Technologies). A 50 μL droplet (containing approximately 300 crypts) was plated in the center of each well of a pre-warmed 24-well plate to form a dome. After polymerization of the Matrigel domes, 750 μL of OGM was added per well. Crypts were cultured at 37 °C in a humidified atmosphere of 5% CO₂, and the medium was replaced every 3 days.

### ChIP assay

Intestinal organoids were isolated from mice and cultured in vitro with either DMSO or 10 μM GW4064. After 48 h of treatment, cells were cross-linked with 1% formaldehyde at room temperature for 10 min, followed by quenching with glycine. Chromatin immunoprecipitation (ChIP) was performed using the SimpleChIP Plus Enzymatic Chromatin IP Kit (Magnetic Beads, Cell Signaling Technology, 9005) according to the manufacturer’s instructions. DNA was then purified using the kit-supplied buffers and column-based purification system. The purified DNA was subjected to qPCR analysis using specifically designed primers.

### Dual luciferase reporter gene experimental

Using the JASPAR database (https://jaspar.elixir.no), three putative FXR binding sites were predicted within the promoter region of the mouse TLR3 gene. Accordingly, three promoter fragments (Frag1, Frag2, and Frag3) were amplified by PCR from mouse spleen genomic DNA. These fragments were subsequently cloned into the pGL4.27 luciferase reporter vector to generate promoter-reporter constructs containing Frag1, Frag2, and Frag3 sequences, respectively.Mouse FXR/RXR overexpression plasmids, a Renilla luciferase control vector (pRL-luciferase, Promega), and the luciferase reporter constructs were co-transfected into HEK293T cells. After 24 h of transfection, cells were harvested for analysis. Luciferase activity was measured using the Dual-Luciferase Reporter Gene Assay Kit (Beyotime, RG027) according to the manufacturer’s instructions, and dual-luciferase signals were quantified using Gene5 software.

### Real-time PCR and RNA-seq profiling

Appropriate amounts of mouse tissues stored in liquid nitrogen or freshly cultured *B.caecimuris* were collected for RNA extraction. Total RNA was isolated from frozen tissues or bacterial samples using TRIzol reagent and the standard phenol-chloroform extraction method. cDNA was synthesized from 2 µg of total RNA using a reverse transcription kit. Primer sequences for genes expressed in mouse tissues and *B.caecimuris* are listed in Supplementary Table S3.RT-qPCR was performed using Hieff UNICON® Universal Blue qPCR SYBR Green Master Mix (Yeasen Biotechnology, Shanghai, China) on an ABI 7500 Fast Real-Time PCR System (Applied Biosystems, USA). Each 20 μL reaction contained 1× master mix, forward and reverse primers at a final concentration of 0.2 μmol/L, and template at a final concentration of 0.25 μmol/L. The amplification conditions were as follows: initial denaturation at 95 °C for 2 min, followed by 40 cycles of 95 °C for 10 s and 60 °C for 30 s.All reactions were performed in triplicate and included target-specific positive controls and negative controls, in which DEPC-treated water was used instead of template. Relative gene expression levels in mouse samples were normalized to 18S rRNA, whereas relative mRNA expression levels in *B.caecimuris* were normalized to 16S rRNA. Relative expression was calculated using the 2^−ΔΔCt method.

Appropriate amounts of mouse intestinal tissues were collected from liquid nitrogen, and total RNA was extracted using the UNIQ-10 Column TRIzol Total RNA Isolation Kit (Sangon Biotech, Shanghai, China) according to the manufacturer’s instructions. The concentration, purity, and integrity of the extracted RNA were assessed, and qualified samples were submitted to Sangon Biotech (Shanghai, China) for transcriptome sequencing. Sequencing data were generated using the Illumina HiSeq™ platform.Raw sequencing data were first subjected to quality assessment using FastQC. Adapter sequences, low-quality reads, and reads containing excessive unknown bases were removed. After quality control, clean reads were aligned to the mouse reference genome using Bowtie2. Gene expression levels were subsequently quantified using featureCounts, and differential expression analysis was performed using DEGseq. Differentially expressed genes were defined as those with log₂(fold change) > 1 and qValue < 0.05.All downstream analyses and visualization were performed using IGV (Integrative Genomics Viewer) and relevant R packages, including identification of differentially expressed genes, visualization of expression patterns, and functional analyses. The transcriptomic data have been deposited in the GSA database, with accession numbers provided in the corresponding records.

To investigate the gene expression changes of B.caecimuris following PFOS exposure, RNA was extracted from bacterial cells cultured with or without PFOS. RNA sequencing was performed by Sangon Biotech (Shanghai, Co., Ltd.). For data analysis, raw reads were first filtered to remove low-quality sequences and adapters using Trimmomatic (v0.39). Clean reads were then aligned to the B.caecimuris reference genome using Bowtie2 (v2.4.2), and gene-level read counts were quantified with featureCounts (v2.0.1). Differential expression analysis was conducted using DESeq2 (v1.34.0) with an adjusted P-value threshold of < 0.05 and a log2 fold change cutoff of |log2FC| ≥ 1. Functional enrichment analysis of differentially expressed genes was performed using KEGG and GO databases via clusterProfiler (v4.2.2). All statistical tests were two-sided, and multiple testing correction was applied using the Benjamini-Hochberg method.

### Western blot

Tissues were lysed in RIPA buffer containing protease and phosphatase inhibitors and homogenized thoroughly. Equal amounts of protein extracts were separated by sodium dodecyl sulfate-polyacrylamide gel electrophoresis (SDS-PAGE) and subsequently transferred onto polyvinylidene difluoride (PVDF) membranes. The membranes were blocked with Tris-buffered saline containing 5% bovine serum albumin (BSA) at room temperature for 2 h, followed by incubation with primary antibodies against Tlr3, P65, phospho-P65, IRF3, phospho-IRF3, ZO-1, Occludin, or β-actin at 4 °C overnight. After washing, the membranes were incubated with horseradish peroxidase (HRP)-conjugated secondary antibodies.

### Metagenomic sequencing and analysis

After Sample QC, sample DNA for metagenomics was extracted using MagPure Fast Stool DNA KF Kit B (catalog no. MD5115-02B; Magen Biotechnology Co., Ltd.). 500ng of Meta DNA was fragmented by ultrasound on Covaris E220 (Covaris, Brighton, UK), then selected to 300 bp ∼700 bp by magnet beads size-selection. The selected DNA fragments were repaired, then ligated with indexed adaptor. The ligation product was amplified by PCR, hybridized with exon probe, and captured by streptavidin beads, The Captured DNA was amplified again by PCR, and circularized to get single-stranded circular(ssCir) library. The ssCir library was then amplified through rolling circle amplification (RCA) to obtain DNA nanoball (DNB). The DNB was then loaded to flowcell and sequenced by DNBSEQ Platform.

The bioinformatics analysis started with raw sequence files. To check the quality of the reads, SOAPnuke filter was used, adapters and low-quality reads were removed. The remaining reads were aligned with bowtie2 and assembled using MEGAHIT. Coding sequences were predicted with Prodigal and clustered into unigenes using CD-HIT. For metagenomic phylogenetic analysis, MetaPhlAn 3.0 was used, and then produced a table containing the detected microbes and their relative abundances. eggNOG mapper was used for functional annotation. Diversity analysis, LEfSe (Linear Discriminant Analysis Effect Size) and Spearman’s correlations analysis was conducted by OmicStudio cloud platform ^[46]^. Visualization was performed using cladograms and bar charts generated via the R package ggplot2 (version 3.2.0)

### Metabolomics analysis

Bile acid profiles were determined as previously described ^[47]^. A total of 184 samples were analyzed in this study, including 92 plasma samples and 92 ileum samples. After the pretreatment of plasma and ileum samples was completed, the samples were analyzed using the Eksigent Ultra Liquid Chromatography 100 system, which is coupled with the AB 5600 Triple TOF system (AB SCIEX). The chromatographic column used was the XBridge Peptide BEH C18 column (3.5μm, 2.1 x 100 mm; Waters Corp.). The column temperature was 40℃, the flow rate was 0.4 mL/min, and the mobile phase consisted of A: water + 0.1% formic acid + 10 mM acetic amine, B: 80% methanol + 20% acetonitrile + 0.1% formic acid. The elution gradient began at 35% B (v/v) for 0.5 min; linearly increased to 60% B over 2.5 min; then to 80% B over 7 min; then to 90% B over 6 min; before finally decreasing linearly to 35% B over 4.5 min and holding for 2.5 min for column re-equilibration. Key parameters for ESI-MS analysis in negative mode on the Triple TOF instrument were set as follows: ion source gases 1 and 2 at 50, curtain gas at 30,source temperature 550 °C, ion spray voltage -4500 V. Scans were acquired from m/z 200-800 (TOF-MS) and m/z 50-800 (auto-MS/MS) with an accumulation time of 19.993 min, collision energy of -45 ± 20 V, and declustering potential of -80 V. The internal standards used were CA-d4, LCA-d4, and UDCA-d4. First, calibration curves for each analyte were constructed using six concentration points. Then, a mixed QC sample (15 bile acid standards in a blank matrix) was analyzed after every 10 experimental samples to monitor instrument performance. Based on five replicate injections, the within-batch RSD for all analytes was below 8%, all of which demonstrated excellent method repeatability.The limits of detection (LOD) and quantification (LOQ) for the bile acids are provided in Supplementary Table S4.

### Statistical analysis

The experimental data are shown as the mean ± SEM values. The normal distribution of the data was determined by the Shapiro-Wilk normality test. For statistical comparisons, Student’s ttest or one-way ANOVA with Tukey’s test was used to compare normally distributed variables. Non-normally distributed data were compared by the Mann-Whitney U test or the Kruskal Wallis test. Spearman’s correlations between changes in microbial species and host bile acid levels were calculated based on species with significant differences between the two groups. BenjaminiHochberg procedure with a cutoff of 0.1 was applied to all Spearman’s correlations. p< 0.05 was considered significant. All the bar plots in this study were generated by GraphPad Prism 9.0 (GraphPad Software, La Jolla, CA, USA). For microbiota analysis, community composition analysis, indicator species analysis, alpha/beta diversity analysis was conducted and visualized via R Studio (version 4.2.1). BSH detection was performed using blastp (v 2.13.0+) with a threshold coverage > 80% and identity > 30%.Continuous variables were assessed for normality using the Shapiro-Wilk test and graphical methods. Normally distributed variables were described as mean ± standard deviation, whereas non-normally distributed variables were described as median and interquartile range. Categorical variables were described as counts and percentages.Associations of serum and fecal PFOS with lipid-related indicators were evaluated using partial Spearman correlation analysis. The analyzed lipid-related indicators included TC, TG, HDL-C, LDL-C, VLDL-C, ApoA1, ApoB, ApoA1/B, Lp(a), and AI. The correlation analyses were adjusted for age, sex, body mass index, smoking status, drinking status, systolic blood pressure, and diastolic blood pressure. Correlation coefficients (r) and corresponding P values were reported.Because GS score was available only in a subset of participants, the associations of serum and fecal PFOS with GS score were analyzed separately among the 82 participants with coronary angiography data. Partial Spearman correlation analysis was also used for GS score, with adjustment for the same covariates. For visualization, PFOS and GS score were transformed into covariate-adjusted residuals, and residual scatter plots were generated. Fitted trend lines and 95% confidence interval boundaries were displayed to illustrate the adjusted associations.All statistical tests were two-sided, and P < 0.05 was considered statistically significant. Statistical analyses were performed using R 4.5.3 software.

## Acknowledgments

We sincerely thank Prof. Changtao Jiang (Peking University) for generously providing the intestinal-specific Fxr knockout (FxrΔIE) mice. The study was supported by the National Key Research and Development Program of China (grant nos. 2025YFA1804503, 2021YFA1301202 to Z. Fang.), the National Natural Science Foundation of China (grant nos. 82473677, 82273676, 92357305 to Z. Fang., grant no. 82470527 to K.Y.C., grant no. 32400963 to Q. Wu), the Young Talent Cultivation Fund of the Endocrinology and Metabolic Diseases Discipline, Tianjin Medical University (grant no 2024XKNFM02 to W.Q.), the Tianjin Key Medical Discipline Construction Project (TJYXZDXK-3-007B) (Grant No.ZXY-ZDSYS2025-4 to Z.X.), the Tianjin Natural Science Foundation (grant no. 25JCYBJC00780 to C.Y.), the National Science and Technology Innovation 2030, Noncommunicable Chronic Diseases-National Science and Technology Major Project (grant no. 2024ZD0524300, 2024ZD0524301 to K.Y.C.), Science and Technology Project of Tianjin Municipal Health Committee (grant no. TJWJ2024RC004 to K.Y.C.), the Key Science and Technology Support Project of Tianjin Science and Technology Bureau (grant no. 24ZXGZSY00130 to K.Y.C.), and the Key Research Project of Tianjin Education Commission (grant no 2024ZD031 to K.Y.C.).

## Author contributions

Z.Z.F., Q.W., and K.Y.C. conceptualized and designed the study. L.H.J., S.H., Z.X., R.X.G., J.H.Z., H.W.L., C.Y., Z.H.Z., F.Y.L., Y.T.A., K.X., Y.T.W., X.S.L., G.Q.Q., C.Y.L., M.Q.Z., Z.X.Z., Y.L., and B.N.L. performed the experiments and analyzed the data. L.H.J., S.H., Z.X., and R.X.G. wrote the manuscript. L.H.J., S.H., Z.X., and R.X.G. contributed equally to this work. All authors edited the manuscript and approved the final manuscript.

## References

[1] Global Burden of Cardiovascular Diseases and Risks 2023 Collaborators. Global, Regional, and National Burden of Cardiovascular Diseases and Risk Factors in 204 Countries and Territories, 1990-2023. Journal of the American College of Cardiology. 2025;86(22):2167–2243.

[2] Tamargo IA, Baek KI, Kim Y, Park C, Jo H. Flow-induced reprogramming of endothelial cells in atherosclerosis. Nature Reviews Cardiology. 2023;20(11):738–753.

[3] Roy P, Orecchioni M, Ley K. How the immune system shapes atherosclerosis: roles of innate and adaptive immunity. Nature Reviews Immunology. 2022;22:251–265.

[4] Münzel T, Sørensen M, Hahad O, et al. The contribution of the exposome to the burden of cardiovascular disease. Nature Reviews Cardiology. 2023;20:651–669.

[5] Chakaroun RM, Olsson LM, Bäckhed F. The potential of tailoring the gut microbiome to prevent and treat cardiometabolic disease. Nature Reviews Cardiology. 2023;20:217–235.

[6] Evich MG, Davis MJB, McCord JP, et al. Per- and polyfluoroalkyl substances in the environment. Science. 2022;375(6580):eabg9065.

[7] Cousins IT, Johansson JH, Salter ME, Sha B, Scheringer M. Outside the safe operating space of a new planetary boundary for per- and polyfluoroalkyl substances (PFAS). Environmental Science & Technology. 2022;56(16):11172–11179.

[8] Brunn H, Arnold G, Körner W, et al. PFAS: forever chemicals—persistent, bioaccumulative and mobile. Environmental Sciences Europe. 2023;35:20.

[9] Haug M, Dunder L, Lind PM, et al. Associations of perfluoroalkyl substances (PFAS) with lipid and lipoprotein profiles. Journal of Exposure Science & Environmental Epidemiology. 2023;33:757–765.

[10] Liu B, Zhu L, Wang M, Sun Q. Associations between per- and polyfluoroalkyl substances exposures and blood lipid levels among adults: a meta-analysis. Environmental Health Perspectives. 2023;131(5):056001.

[11] Dunder L, Salihovic S, Varotsis G, et al. Plasma levels of per-and polyfluoroalkyl substances (PFAS) and cardiovascular disease: results from two independent population-based cohorts and a meta-analysis. Environment International. 2023;181:108250.

[12] Biggeri A, Stoppa G, Facciolo L, et al. All-cause, cardiovascular disease and cancer mortality in the population of a large Italian area contaminated by perfluoroalkyl and polyfluoroalkyl substances (1980-2018). Environmental Health. 2024;23:42.

[13] Zhu L, Liu B, Hu Y, et al. Per- and polyfluoroalkyl substances, apolipoproteins and the risk of coronary heart disease in US men and women. Environmental Health. 2024;23:108.

[14] Wang D, Tan Z, Yang J, et al. Perfluorooctane sulfonate promotes atherosclerosis by modulating M1 polarization of macrophages through the NF-κB pathway. Ecotoxicology and Environmental Safety. 2023;249:114384.

[15] Violi F, Cammisotto V, Bartimoccia S, et al. Gut-derived low-grade endotoxaemia, atherothrombosis and cardiovascular disease. Nature Reviews Cardiology. 2023;20:24–37.

[16] Collins SL, Stine JG, Bisanz JE, et al. Bile acids and the gut microbiota: metabolic interactions and impacts on disease. Nature Reviews Microbiology. 2023;21:236–247.

[17] Birkeland E, Gharagozlian S, Valeur J, et al. Short-chain fatty acids as a link between diet and cardiometabolic risk: a narrative review. Lipids in Health and Disease. 2023;22:40.

[18] Wang B, Qiu J, Lian J, Yang X, Zhou J. Gut metabolite trimethylamine-N-oxide in atherosclerosis: from mechanism to therapy. Frontiers in Cardiovascular Medicine. 2021;8:723886.

[19] Zhang L, Rimal B, Nichols RG, et al. Perfluorooctane sulfonate alters gut microbiota-host metabolic homeostasis in mice. Toxicology. 2020;431:152365.

[20] Chen Q, Sun S, Mei C, et al. Capabilities of bio-binding, antioxidant and intestinal environmental repair jointly determine the ability of lactic acid bacteria to mitigate perfluorooctane sulfonate toxicity. Environment International. 2022;166:107388.

[21] De Filippis F, Valentino V, Sequino G, et al. Exposure to environmental pollutants selects for xenobiotic-degrading functions in the human gut microbiome. Nature Communications. 2024;15:4482.

[22] Lindell AE, Grießhammer A, Michaelis L, et al. Human gut bacteria bioaccumulate per- and polyfluoroalkyl substances. Nature Microbiology. 2025;10:1630–1647.

[23] Yan, Y., Lei, Y., et al. Bacteroides uniformis-induced perturbations in colonic microbiota and bile acid levels inhibit TH17 differentiation and ameliorate colitis developments. NPJ Biofilms Microbiomes. 2023 Aug 14;9(1):56.

[24] Zhao Q, Dai MY, Huang RY, et al. Parabacteroides distasonis ameliorates hepatic fibrosis potentially via modulating intestinal bile acid metabolism and hepatocyte pyroptosis in male mice. Nat Commun. 2023 Apr 1;14(1):1829.

[25] Chen Z, Chen H, Huang W, et al. *Bacteroides fragilis* alleviates necrotizing enterocolitis through restoring bile acid metabolism balance using bile salt hydrolase and inhibiting FXR-NLRP3 signaling pathway. Gut Microbes. 2024 Jan-Dec;16(1):2379566.

[26] Qi Z, Zhang W, Zhang P, et al. The gut microbiota-bile acid-TGR5 axis orchestrates platelet activation and atherothrombosis. Nat Cardiovasc Res. 2025 May;4(5):584–601.

[27] Ko S, Anzai A, Liu X, et al. Social Bonds Retain Oxytocin-Mediated Brain-Liver Axis to Retard Atherosclerosis. Circ Res. 2025 Jan 3;136(1):78–90.

[28] Sun L, Xie C, Wang G, et al. Gut microbiota and intestinal FXR mediate the clinical benefits of metformin. Nature Medicine. 2018;24:1919–1929.

[29] Jang S. AcrAB–TolC, a major efflux pump in Gram-negative bacteria: toward understanding its operation mechanism. BMB Reports. 2023;56:326–334.

[30] Nikaido H, Pagès JM. Broad-specificity efflux pumps and their role in multidrug resistance of Gram-negative bacteria. FEMS Microbiology Reviews. 2012;36:340–363.

[31] Blanco P, Hernando-Amado S, Reales-Calderón JA, et al. Bacterial multidrug efflux pumps: much more than antibiotic resistance determinants. Microorganisms. 2016;4:14.

[32] Rimal B, Collins SL, Tanes CE, et al. Bile salt hydrolase catalyses formation of amine-conjugated bile acids. Nature. 2024;626:859–863.

[33] Matsumoto M, Seya T. TLR3: interferon induction by double-stranded RNA including poly(I:C). Advanced Drug Delivery Reviews. 2008;60:805–812.

[34] Vercammen E, Staal J, Beyaert R. Sensing of viral infection and activation of innate immunity by Toll-like receptor 3. Clinical Microbiology Reviews. 2008;21:13–25.

[35] Claudel T, Staels B, Kuipers F. The farnesoid X receptor: a molecular link between bile acid and lipid and glucose metabolism. Arteriosclerosis, Thrombosis, and Vascular Biology. 2005;25:2020–2030.

[36] Massafra V, Pellicciari R, Gioiello A, van Mil SWC. Progress and challenges of selective farnesoid X receptor modulation. Pharmacology & Therapeutics. 2018;191:162–177.

[37] Wu Q, Sun L, Hu X, et al. Suppressing the intestinal farnesoid X receptor/sphingomyelin phosphodiesterase 3 axis decreases atherosclerosis. Journal of Clinical Investigation. 2021;131:e142865.

[38] Bile acid receptor FXR promotes intestinal epithelial ferroptosis and subsequent ILC3 dysfunction in neonatal necrotizing enterocolitis. Immunity. 2025.

[39] Leonor GB, E. CL. Streamlined Genetic Manipulation of Diverse Bacteroides and Parabacteroides Isolates from the Human Gut Microbiota. MBio. 2019;10(4):10.1128/mbio.01762-19.

[40] Minoru I, Haruyuki NI, Shin W, et al. Efficient Electrotransformation of Bacteroides fragilis. Appl Environ Microbiol. 2010;76(10):3325–3332.

[41] Jiang L, Hong Y, et al. The Role of Fecal Microbiota in Liver Toxicity Induced by Perfluorooctane Sulfonate in Male and Female Mice. Environ Health Perspect. 2022 Jun;130(6):67009.

[42] Wang Z, Yao J, et al. Comparative Hepatotoxicity of a Novel Perfluoroalkyl Ether Sulfonic Acid, Nafion Byproduct 2 (H-PFMO2OSA), and Legacy Perfluorooctane Sulfonate (PFOS) in Adult Male Mice. Environ Sci Technol. 2022 Jul 19;56(14):10183–10192.

[43] Zhang Y, Wang X, Lin J, Liu J, et al. A microbial metabolite inhibits the HIF-2α-ceramide pathway to mediate the beneficial effects of time-restricted feeding on MASH. Cell Metab. 2024 Aug 6;36(8):1823–1838.e6.

[44] She J, Tuerhongjiang G, Guo M, Liu J, et al. Statins aggravate insulin resistance through reduced blood glucagon-like peptide-1 levels in a microbiota-dependent manner. Cell Metab. 2024 Feb 6;36(2):408–421.e5.

[45] Bauer PV, Duca FA, Waise TMZ, et al. Lactobacillus gasseri in the Upper Small Intestine Impacts an ACSL3-Dependent Fatty Acid-Sensing Pathway Regulating Whole-Body Glucose Homeostasis. Cell Metab. 2018 Mar 6;27(3):572–587.e6.

[46] Qin G, Xu Z, Yuan C, et al. 4-Hydroxyphenanthrene exacerbates obesity by altering gut microbiota and bile acid metabolism. Nat Commun. 2026 Mar 13.

[47] Wu Q, Liang X, Wang K, et al. Intestinal hypoxia-inducible factor 2α regulates lactate levels to shape the gut microbiome and alter thermogenesis. Cell Metab. 2021 Oct 5;33(10):1988–2003.e7.

